# Genetic Analysis of The Endangered Cleveland Bay Horse: A Century of Breeding Characterised by Pedigree and Microsatellite Data

**DOI:** 10.1101/2020.05.19.104182

**Authors:** Andrew Dell, Mark Curry, Kelly Yarnell, Gareth Starbuck, Philippe B. Wilson

## Abstract

The Cleveland Bay horse is one of the oldest equines in the United Kingdom, with pedigree data going back almost 300 years. The studbook is essentially closed and because of this, there are concerns about loss of genetic variation across generations. The breed is one of five equine breeds listed as “critical” (<300 registered adult breeding females) by the UK Rare Breeds Survival Trust in their annual Watchlist. Due to their critically endangered status, the current breadth of their genetic diversity is of concern, and assessment of this can lead to improved breed management strategies. Herein, both genealogical and molecular methods are combined in order to assess founder representation, lineage, and allelic diversity. Data from 16 microsatellite loci from a reference population of 402 individuals determined a loss of 91% and 48% of stallion and dam lines, respectively in the reference population. Only 3 ancestors determine 50% of the genome in the living population, with 70% of maternal lineage being derived from 3 founder females, and all paternal lineages traced back to a single founder stallion. Methods and theory are described in detail in order to demonstrate the scope of this analysis for wider conservation strategies. We quantitatively demonstrate the critical nature of the genetic resources within the breed, and offer a perspective on implementing this data in considered breed management strategies.

## Introduction

In recent years there has been substantial interest in quantifying the genetic diversity of equine breeds using pedigree, [1] molecular data [2] or a combination of both sources [3] in order to implement effective breed management strategies. The effectiveness of the use of both data types in the understanding and management of rare and native equine breeds have been investigated using both theoretical modelling, and studies of closed studbooks.

The Cleveland Bay horse is a heritage British breed which has its origins in the Cleveland Hills of Northern England [4]. The first studbook was published in 1885, and this contains retrospective pedigrees of animals dating back to 1732 providing a closed non-Thoroughbred studbook dating back almost 300 years and for more than 38 generations. In addition, the breed Society now has a mandatory policy of microsatellite-based parentage testing at the time of registration. Unrestricted access to the microsatellite test data, as well as the stud book records provides a rare opportunity to evaluate both methods of assessing genetic diversity within the breed and, in turn, provides comprehensive guidance to breeders in terms of conservation practice for this endangered breed [5], whilst providing an important and potentially wide-ranging tool for wider conservation practices both *in situ* and *ex situ in vivo*.

The Cleveland Bay is a warm-blooded equine; a product of a cross of hot-blooded Oriental / Barb /Turkish or Mediterranean stock on the cold-blooded Northern European heavy draught horse [6]. It is reputed to have evolved in the matriline from the now extinct Chapman horse, which early records show were being bred on the monastic estates of the region well before the dissolution of the monasteries in the mid 16th century [4].

Although stated as being “free of blood” in the first three volumes of the studbook,[7] early research into the founders of the breed recognised the contribution on the male side by some notable Thoroughbred stallions that were standing at stud or travelling in the region in the late 18th and early 19th Centuries [8].

Over the years the breed has been used extensively as both a work horse and a riding horse, and has been crossed with other breeds to produce carriage horses [9]. Indeed, at one time there was a separate breed society with its own studbook – The Yorkshire Coach Horse Society – for such animals [10]. It is reputed that in the United States, Buffallo Bill used Cleveland Bay horses in his “Wild West Show” [11]. Such has been the desirability of the pure Cleveland Bay for contributing weight carrying capacity when crossed with other equine breeds, that they have been exported globally [9]. In addition to North America, the breed has been exported to Australasia, Pakistan and Japan; a Cleveland Bay stallion stands at the Imperial stud [8].

The fashion for such effective cross-bred horses is one factor that brought the pure-bred Cleveland Bay horse to the edge of extinction. The substantial decrease in population size of the breed following the First World War when large numbers of Cleveland Bay horses were used to haul artillery on the battlefields of Northern Europe led to sustainability concerns regarding the remaining genetic resources of the breed [9]. The popularity of the breed continued to decline in the 1920s and 30s as the increasing use of motorised transport reduced the need for carriage horses. Moreover, following the technological developments of the Second World War, further mechanisation was implemented in farming practice and the purpose of the Cleveland Bay was further diminished [12].

In an attempt to improve the diversity of the home-based breeding population, the stallion Farnley Exchange was brought back from the United States of America (USA) in 1945 to stand at stud [9]. By the early 1960s there were only four stallions of breeding age left in existence and the breed is known to have gone through a genetic bottleneck at this time [8].

In the 1960s HM the Queen purchased the stallion Mulgrave Supreme, thus preventing his export, and stood him at public stud, both to promote, and help conserve the genetic diversity of the breed in the United Kingdom. Since that time the breed has seen a moderate recovery in numbers, partly because of patronage of the breed society by HM the Queen and the use of Cleveland Bay horses at the Royal Mews.

By the late 1990s, between 35 and 50 pure bred animals were being registered annually by the Cleveland Bay Horse Society (CBHS), whose studbook now includes animals being bred both in the United Kingdom, Europe, North America and Australasia [13].

The breed is one of only five equines listed as “Critical” by the UK Rare Breeds Survival Trust, indicating that the population has less than 300 breeding females. Earlier investigation of the CBHS Studbook records [7] indicated there were eight female ancestry lines existing within the breed.

A more recent study [13] restricted to animals entered in the CBHS studbook between 1934 and 1995, highlighted the limited genetic diversity in the breed and the increasing levels of inbreeding. It was recognised that further in-depth analysis of the status of the breed would be needed in order to aid in the development of breed management plans.

Herein, we develop a comparative analysis of the genetic diversity in the Cleveland Bay Horse population using both genealogical and molecular methods and provide recommendations in order to support a global breed conservation strategy for the Cleveland Bay Horse, whilst sequentially detailing the theory and practice inherent in our approach leading to its applicability in the conservation of endangered breeds and species *in vivo*.

## Results and Discussion

### Pedigree Completeness

The pedigree file included a total of 5422 animals, of which 2661 were male and 2761 were female. The reference population of 402 individual animals consisted of 193 male and 209 females.

The pedigree file was analysed to assess the number of fully traced generations for each individual, the maximum number of generations traced and the equivalent complete generations for each animal. The maximum number of traced generations was 36. Percentage average population completeness for each year of birth considering 1 through 6 generations are shown in Figure 1.

**Figure 1.**
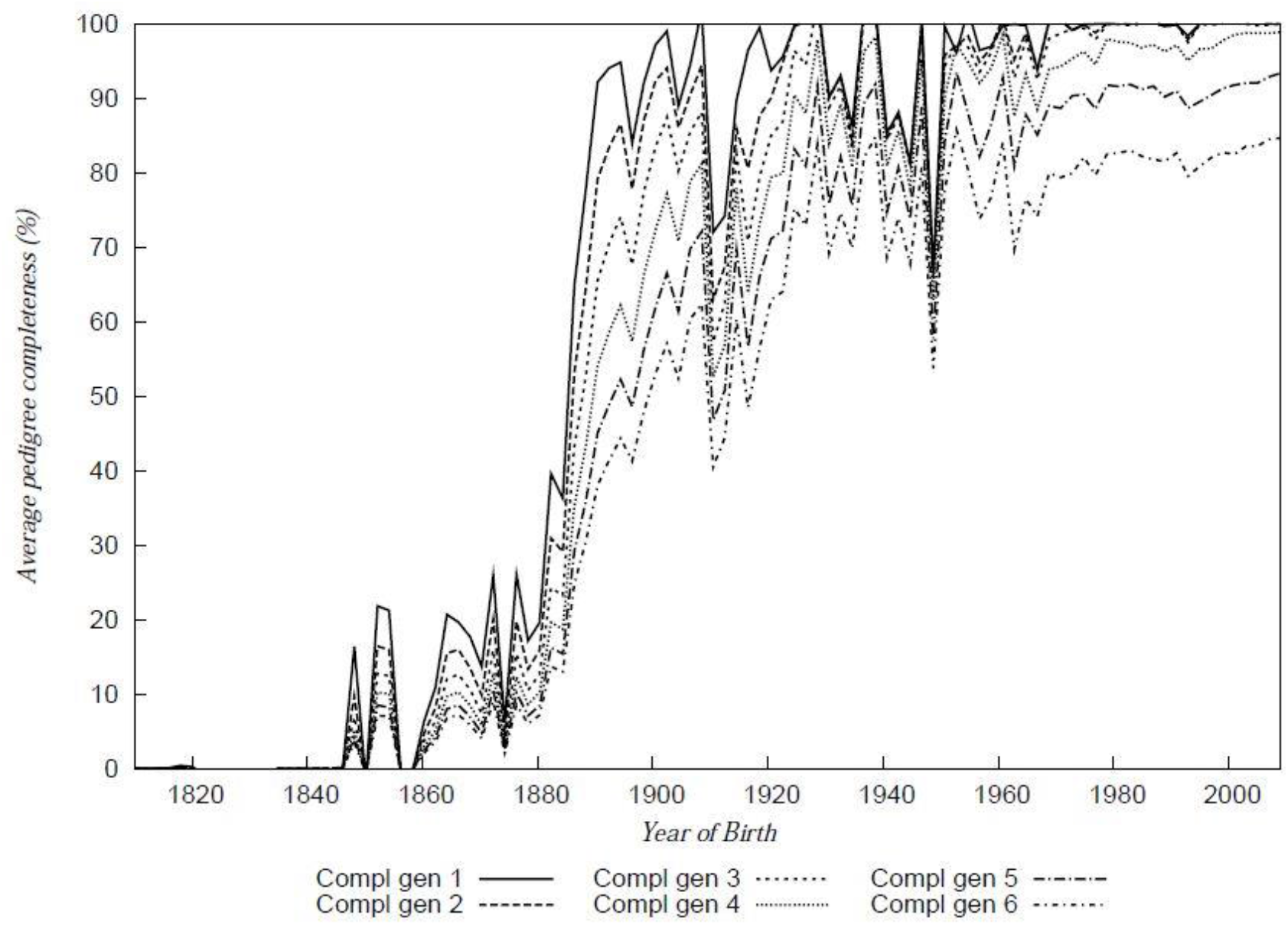
Percentage pedigree completeness over 6 generations. Average percentage completeness (%) is shown as a factor of individual birth year.

Percentage population completeness for the reference population up to 6 generations is shown in Table 1.

**Table 1.** Pedigree Completeness over 6 generations estimated from breed society records and pedigree recording data.

| Generations | Completeness (%) |
| --- | --- |
| 1 | 100 |
| 2 | 100 |
| 3 | 99.9 |
| 4 | 98.6 |
| 5 | 92.6 |
| 6 | 83.7 |

### Average Generation Interval

The average generation interval for each breeding year is shown in Figure 2. This was found to range between 5.5 and 13 years, being at a minimum in the immediate post WW2 period 1946 to 1950, which coincides with the genetic bottleneck previously identified by Walling (1994).

**Table 2.** Average Generation Interval by pathway.

| Pathway | Average generation interval ( years) |
| --- | --- |
| Sire son | 9.2 |
| Sire daughter | 10.0 |
| Dam son | 9.6 |
| Dam daughter | 9.3 |
| Whole pop | 9.6 |

**Figure 2.**
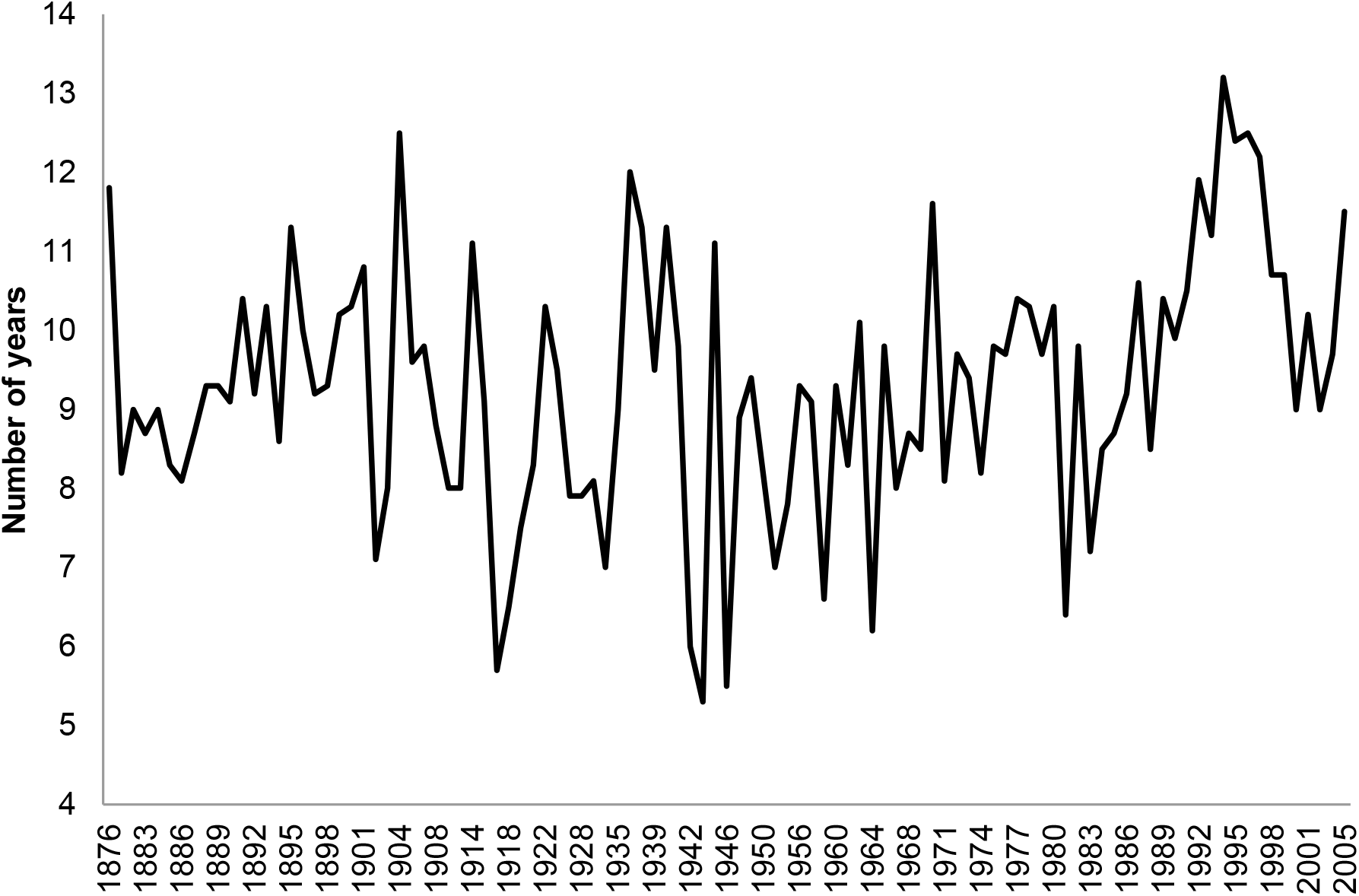
Average Generation interval for whole population calculated as the average age of parents at the birth of offspring which in turn produce the next generation of breeding individuals.

### Founder and Ancestor representation

A total of 11 stallion lines were identified in the pedigree. A single paternal ancestry line is present in the reference (living) population.

Analysis of the female members of the studbook identified a total of 17 dam lines. Nine of these maternal ancestry lines are present in the reference population. Three of these lines (2,4 & 9) are only represented, in direct female descent, by either a single individual or two individual animals (Table 3). The three most common maternal lines constitute 70% of the present female population. However, analysis of the relative contributions of the most influential maternal ancestry lines to the genome of the reference population reveals that some of the lines least well represented in direct descent in fact continue to make a substantial genetic contribution as shown in Table 3.

**Table 3.**
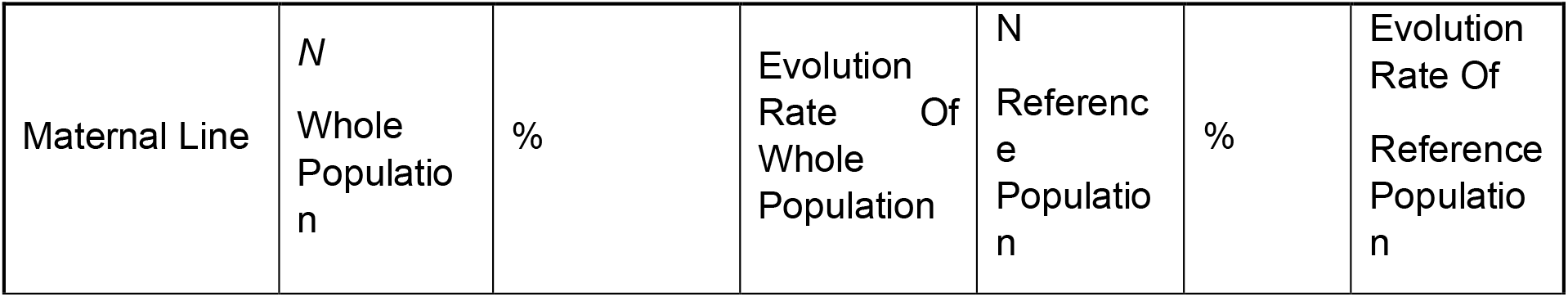
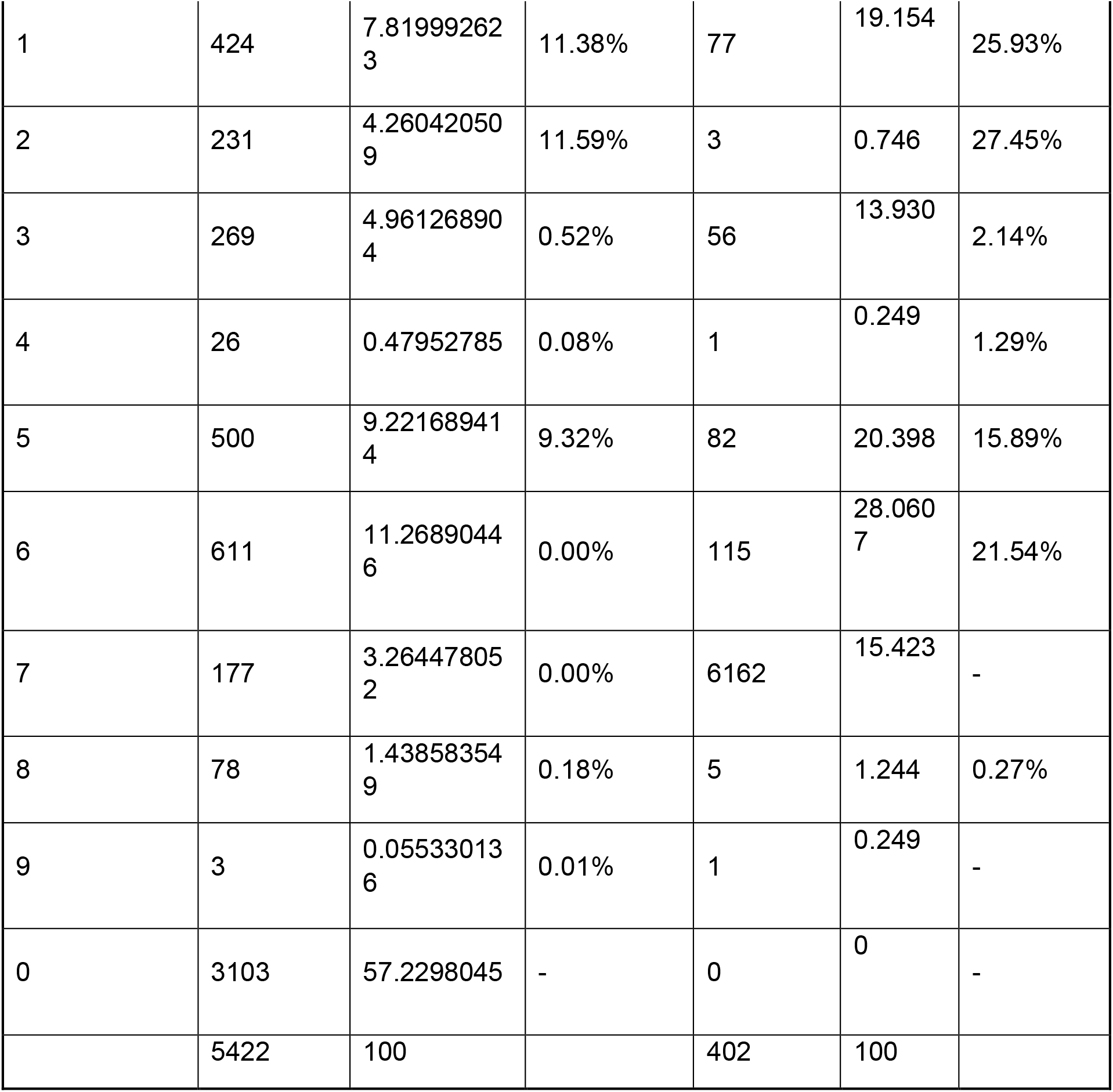
Relative contributions of maternal ancestry lines to the evolution of the whole and reference (1997-2006) populations.

| Maternal Line | N<br>Whole<br>Populatio<br>n | % | Evolution<br>Rate<br>Of<br>Whole<br>Population | N<br>Referenc<br>e<br>Populatio<br>n | % | Evolution<br>Rate Of<br>Reference<br>Populatio<br>n |
| --- | --- | --- | --- | --- | --- | --- |
| 1 | 424 | 7.81999262<br>3 | 11.38% | 77 | 19.154 | 25.93% |
| 2 | 231 | 4.26042050<br>9 | 11.59% | 3 | 0.746 | 27.45% |
| 3 | 269 | 4.96126890<br>4 | 0.52% | 56 | 13.930 | 2.14% |
| 4 | 26 | 0.47952785 | 0.08% | 1 | 0.249 | 1.29% |
| 5 | 500 | 9.22168941<br>4 | 9.32% | 82 | 20.398 | 15.89% |
| 6 | 611 | 11.2689044<br>6 | 0.00% | 115 | 28.060<br>7 | 21.54% |
| 7 | 177 | 3.26447805<br>2 | 0.00% | 6162 | 15.423 | - |
| 8 | 78 | 1.43858354<br>9 | 0.18% | 5 | 1.244 | 0.27% |
| 9 | 3 | 0.05533013<br>6 | 0.01% | 1 | 0.249 | - |
| 0 | 3103 | 57.2298045 | - | 0 | 0 | - |
|  | 5422 | 100 |  | 402 | 100 |  |

Analysis using GENES [14] identified 194 founders in total of which 28 were represented in the reference population. The mean retention was 0.035. The number of founder genomes surviving was 6.285. Calculations on the same population using CFC [15] show the founder genome equivalent to be 2.366 with the effective number of non-founders only 2.379. The proportion of ancestry known was 0.330 reflecting the fact that in early volumes of the studbook only a record of the sire of an individual animal was made. The Number of Ancestors contributing to the population was 424 and the number of ancestors describing 50% of the genome was 7.

The number of Ancestors contributing to the Reference Population was calculated as 31. The Effective Number of Founders/Ancestors [16] for the Reference Population were 40 and 9, respectively. The number of ancestors describing 50% of the genome of the living population was 3. Ancestors were selected following Boichard *et al.* (1997), while founders were selected by their individual Average Relatedness coefficient (AR).

### Inbreeding Analysis

Across the whole analysed dataset, *F*= 7.8% with an associated mean average relatedness of 8.3%. Figure 3 shows Inbreeding and additive relationship coefficients by birth year between 1900 to 2006.

**Figure 3.**
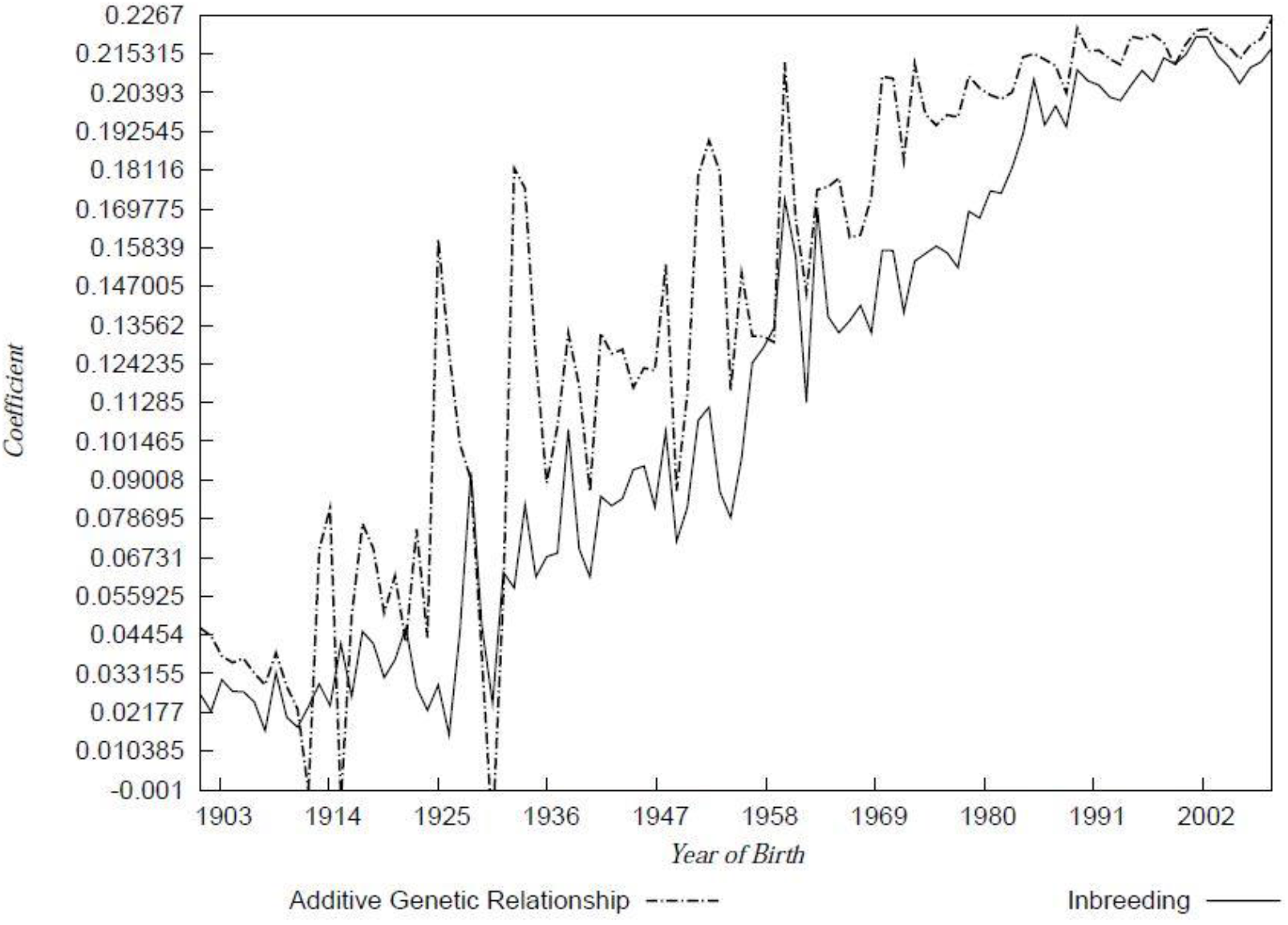
Inbreeding Coefficient and Additive Genetic Relationship 1900 to 2006 as a function of birth year of individuals.

**Table 4.**
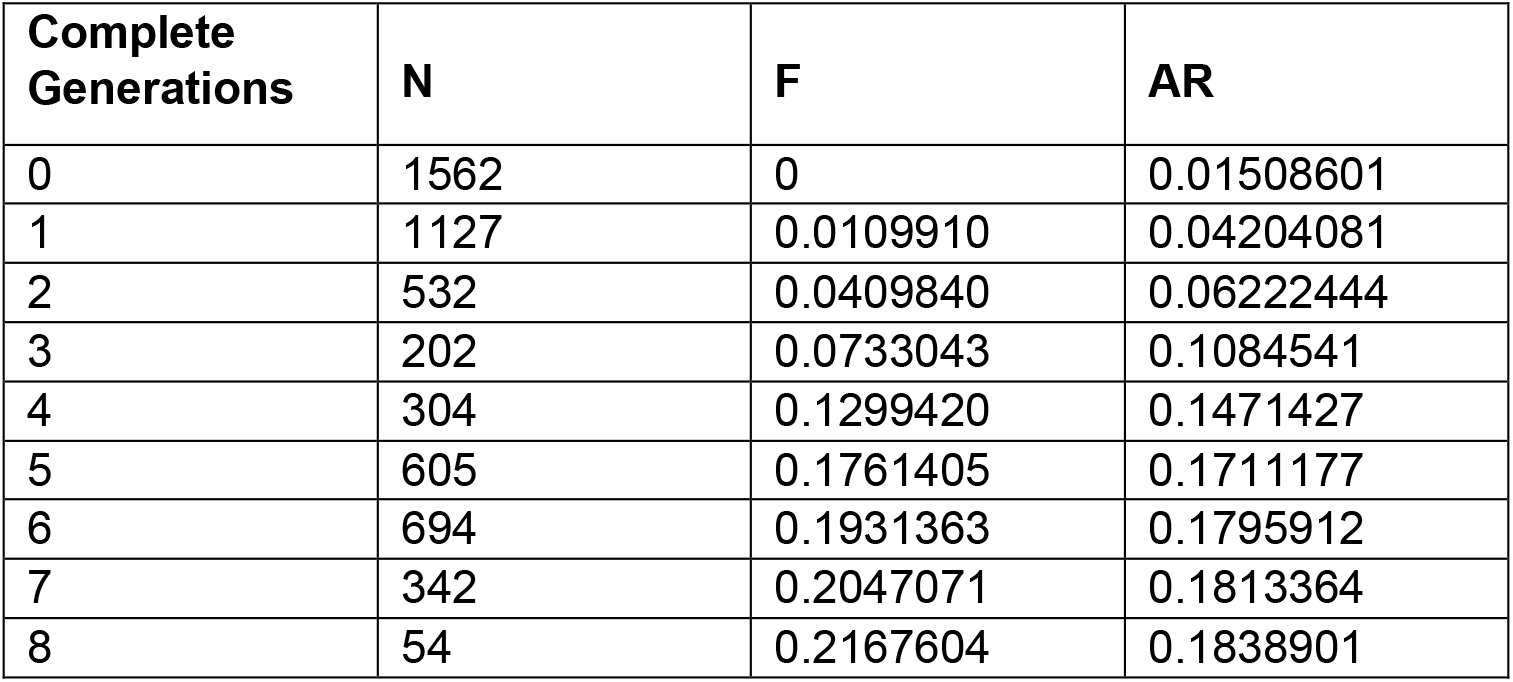
Change in inbreeding coefficient and average relatedness for 7 fully traced generations.

The average rate of change of the additive genetic relationships between 1901 and 2009 for the Cleveland Bay Horse breed was 0.00202 per year based on the slope regression. This results in a *Δf* per generation of 0.02629. The rate of change of the average inbreeding coefficients based on slope regression between 1901 and 2009 was 0.00214, which represents a *ΔF* per generation of 0.02709. The effective population sizes for the Cleveland Bay Horse breed, based on *Δf* and *ΔF* were 19 and 18, respectively. The pattern of inbreeding over the period 1997 to 2006 during which the reference population was foaled is shown in Table 5, with data calculated using POPREP [17].

**Table 5.** Inbreeding Coefficients, *F,* of reference population animals by birth year 1997-2006.

| Year | Number of Animals | Minimum Inbreeding | Maximum Inbreeding | Average | Std |
| --- | --- | --- | --- | --- | --- |
| 1997 | 57 | 0.1327 | 0.2943 | 0.2072 | 0.0305 |
| 1998 | 46 | 0.1540 | 0.2943 | 0.2139 | 0.0254 |
| 1999 | 54 | 0.1448 | 0.3156 | 0.2126 | 0.0317 |
| 2000 | 64 | 0.1654 | 0.3079 | 0.2139 | 0.0269 |
| 2001 | 37 | 0.1783 | 0.3132 | 0.2186 | 0.0250 |
| 2002 | 46 | 0.1830 | 0.3084 | 0.2218 | 0.0263 |
| 2003 | 52 | 0.1830 | 0.3017 | 0.2173 | 0.0227 |
| 2004 | 63 | 0.1629 | 0.2852 | 0.2133 | 0.0231 |
| 2005 | 54 | 0.1100 | 0.2580 | 0.2102 | 0.0219 |
| 2006 | 76 | 0.0925 | 0.2616 | 0.2065 | 0.0253 |

### Effective population size

Effective population size was calculated based on both the rate of inbreeding and the number of parents. The results are tabulated in Table 6 for the period 1997 to 2006.

**Figure 4.**
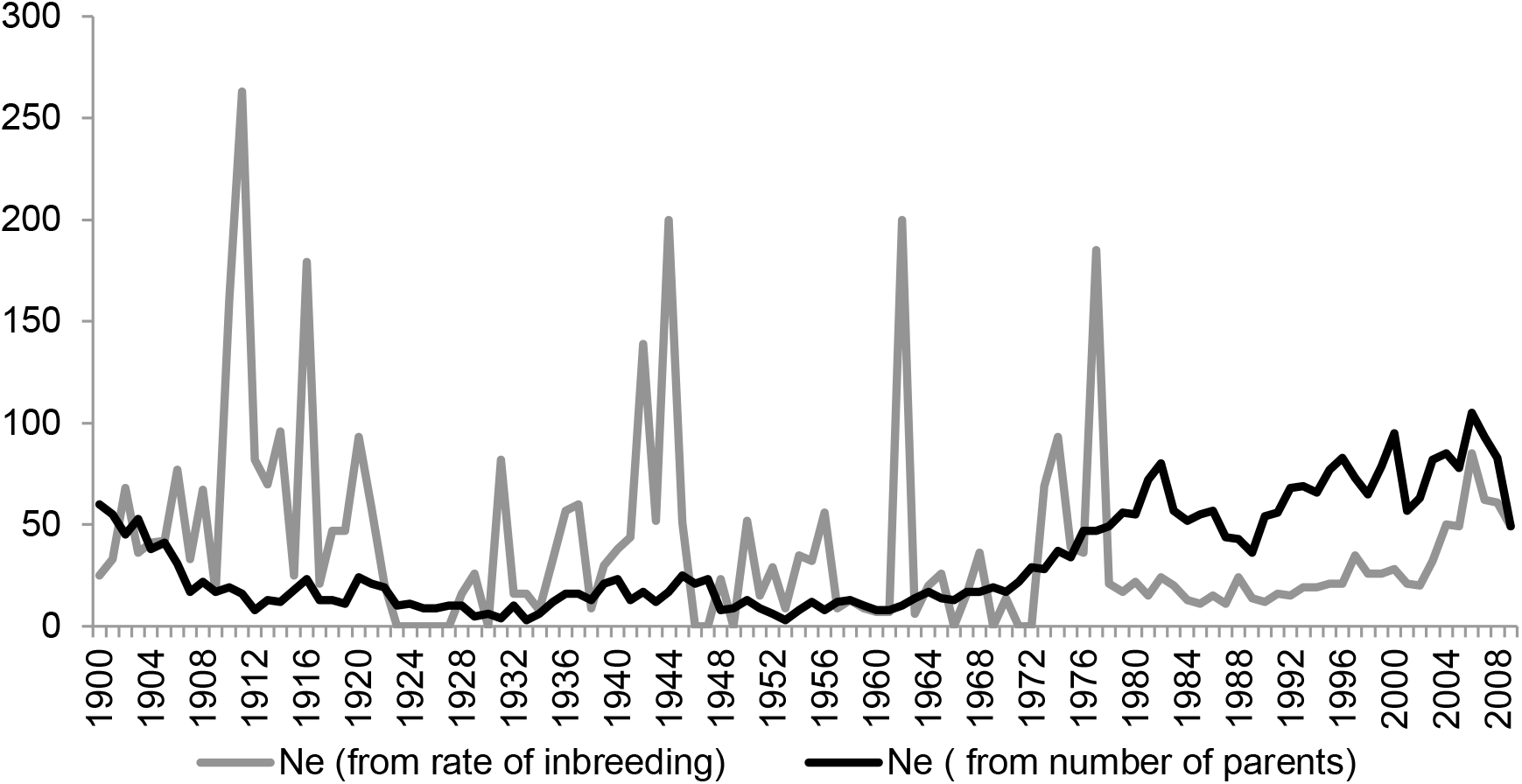
Effective Population Size from rate of change of inbreeding (grey series), and number of parents (black series) calculated with POPREP [17].

**Table 6.**
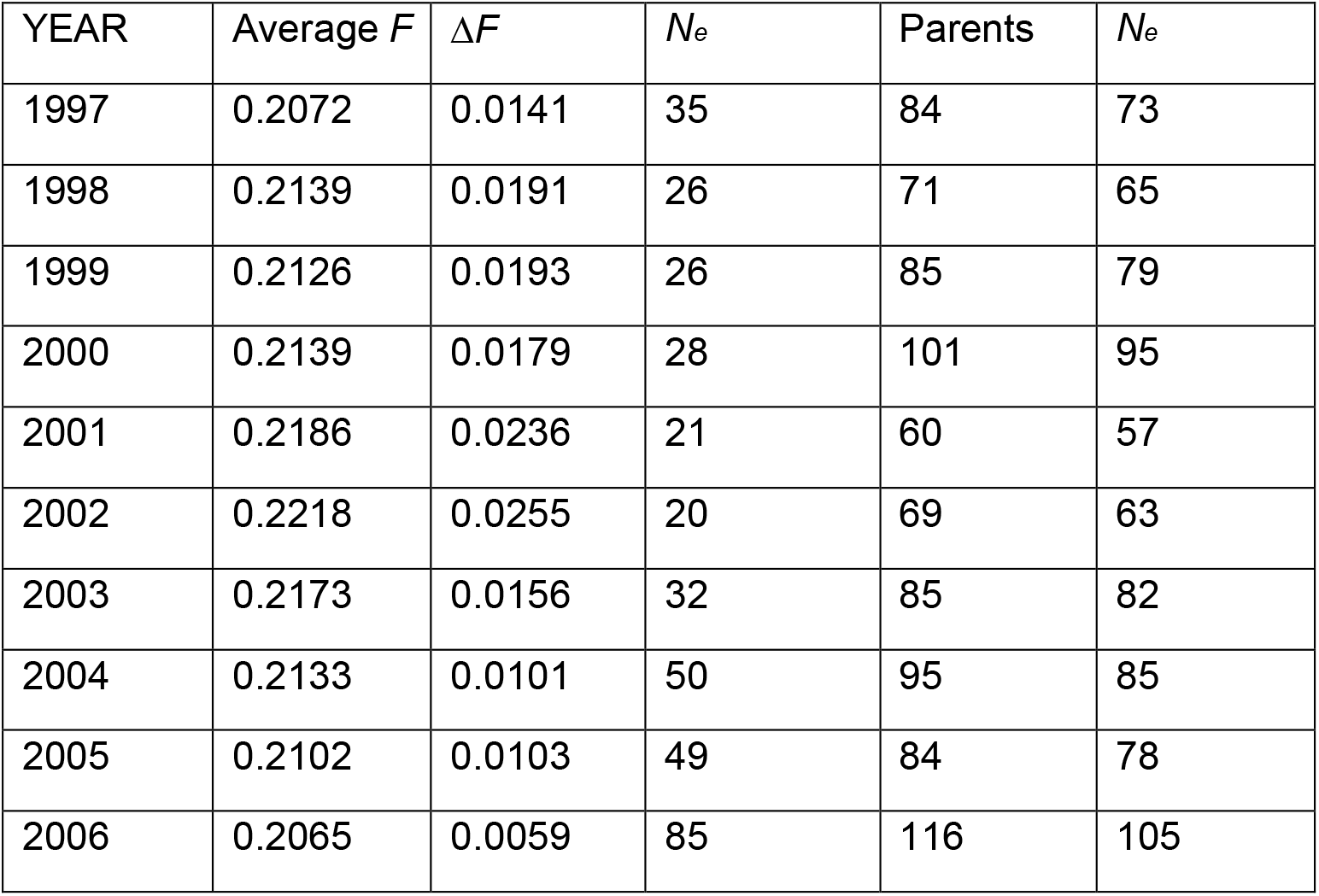
Effective Population Size (*N*_*e*_) in Reference Population.

### Microsatellite Variation

The total number of alleles found for 16 microsatellite loci within the reference population was 99. The mean number of alleles per locus was 6.19 ranging from 4 to 10. The mean Observed Heterozygosity (*H*_*o*_) ranged between 0.052(HTG7) and 0.716 (VHL20) the mean being 0.0.4486 whilst the mean Expected Heterozygosity (*H*_*e*_) was 0.5341. The mean Polymorphic Information Content (PIC) was 0.4957. The highest values for *H*_*e*_ and PIC were found for microsatellite LEX33 whilst the lowest were found for microsatellite HTG6 (Table 7).

**Table 7.** Summary statistics for the 16 microsatellite loci analysed. N_a_ represents the number of alleles; *N*, the sample size; *H*_*o*_ the Observed Heterozygosity; *H*_*e*_ the Expected Heterozygosity; *PIC* the Polymorphic Information Content; HW the departure from Hardy-Weinberg equilibrium; *F* the Fixation Index; and *N*_*m\**_ the Gene flow estimated from *F*_ST_ = 0.25(1 - *F*_*ST*_)/*F*_*ST*_.

| Locus | $N_a$ | $N$ | $H_o$ | $H_e$ | PIC | HW | $F$ | $F_{IS}$ | $F_{IT}$ | $F_{ST}$ | $N_{m^*}$ |
| --- | --- | --- | --- | --- | --- | --- | --- | --- | --- | --- | --- |
| VHL20 | 5 | 402 | 0.716 | 0.697 | 0.640 | NS | - | -0.290 | - | 0.113 | 1.968 |
| HTG4 | 4 | 402 | 0.463 | 0.434 | 0.378 | NS | - | -0.105 | 0.0 | 0.146 | 1.463 |
| AHT4 | 7 | 402 | 0.530 | 0.529 | 0.507 | NS | - | -0.268 | - | 0.121 | 1.810 |
| HMS7 | 5 | 402 | 0.697 | 0.706 | 0.655 | ND | +0.0 | -0.293 | - | 0.110 | 2.017 |
| HTG6 | 5 | 402 | 0.067 | 0.173 | 0.168 | NS | +0.5 | -0.240 | 0.3 | 0.510 | 0.240 |
| AHT5 | 7 | 402 | 0.669 | 0.684 | 0.629 | NS | +0.0 | -0.345 | - | 0.134 | 1.622 |
| HMS6 | 5 | 402 | 0.590 | 0.572 | 0.488 | NS | - | -0.271 | 0.0 | 0.245 | 0.769 |
| ASB2 | 8 | 402 | 0.550 | 0.578 | 0.528 | NS | +0.0 | -0.219 | - | 0.169 | 1.227 |
| HTG10 | 4 | 402 | 0.687 | 0.675 | 0.613 | NS | - | -0.070 | 0.1 | 0.174 | 1.189 |
| HTG7 | 4 | 402 | 0.052 | 0.180 | 0.172 | ND | +0.6 | -0.252 | 0.5 | 0.614 | 0.157 |
| HMS3 | 7 | 402 | 0.187 | 0.203 | 0.196 | ND | +0.0 | -0.197 | 0.0 | 0.199 | 1.007 |
| HMS2 | 6 | 402 | 0.057 | 0.176 | 0.169 | ND | +0.5 | -0.243 | 0.4 | 0.559 | 0.197 |
| ASB17 | 10 | 402 | 0.500 | 0.780 | 0.744 | ND | +0.2 | -0.020 | 0.3 | 0.329 | 0.509 |
| ASB23 | 6 | 402 | 0.639 | 0.759 | 0.722 | *** | +0.0 | -0.082 | 0.0 | 0.155 | 1.362 |
| LEX3 | 6 | 402 | 0.201 | 0.591 | 0.548 | ND | +0.4 | 0.444 | 0.5 | 0.119 | 1.858 |
| LEX33 | 10 | 402 | 0.575 | 0.805 | 0.773 |  | +0.1 | -0.196 | 0.0 | 0.189 | 1.073 |
| MEAN | 6.1 | 402 | 0.448 | 0.534 | 0.495 |  |  | -0.166 | 0.0 | 0.243 | 1.154 |

Significant deviations from HWE were observed for microsatellites AHT4, HTG1, LEX3 and LEX33.

**Table 8.** Summary of the microsatellite analysis results on a subpopulation by matriline basis and for the full dataset, where MNA represents the mean number of alleles per locus.

| SUBPOPULATION | *N* | Observed Heterozygosity | Expected Heterozygosity | Unbiased Expected Heterozygosity | PIC | MNA | Polymorphic Loci |
| --- | --- | --- | --- | --- | --- | --- | --- |
| 1 | 77 | 0.5590 | 0.5713 | 0.5769 | 0.4051 | 4.9375 | 100.00% |
| 2 | 3 | 0.4573 | 0.3196 | 0.3937 | 0.1939 | 1.8125 | 62.50% |
| 3 | 56 | 0.5893 | 0.6078 | 0.5978 | 0.4198 | 4.5625 | 100.00% |
| 4 | 1 | 0.5833 | 0.4208 | 0.5222 | 0.2860 | 1.9333 | 62.50% |
| 5 | 82 | 0.5919 | 0.6217 | 0.6009 | 0.4262 | 4.5 | 100.00% |
| 6 | 115 | 0.5659 | 0.6092 | 0.5749 | 0.4152 | 4.875 | 100.00% |
| 7 | 61 | 0.5818 | 0.6138 | 0.5857 | 0.4199 | 4.6875 | 100.00% |
| 8 | 4 | 0.6258 | 0.5483 | 0.5626 | 0.3499 | 3.0625 | 93.75% |
| GRADING | 3 | 0.4722 | 0.3251 | 0.4021 | 0.1510 | 1.75 | 62.50% |
| MEAN |  | 0.5585 | 0.5153 | 0.5352 | 0.3408 | 3.568978 | 86.81% |

Across the reference population there is complete heterozygosity. However, at subpopulation level 3, groups show homozygosity at multiple loci. Female Line 2 is 62.5% polymorphic with fixation at HMS3 and LEX3. Female Line 4 is 62.5% polymorphic with fixation of alleles at HMS3, ASB23, HTG4, HTG10 and LEX3. Female Line 8 is 93.75% polymorphic with fixation at LEX3.

Allele frequencies are more restricted in populations 2, 4 and nine (Figure 5), as is the expected heterozygosity. This will be influenced by the smaller membership and corresponding sample size for these subpopulations.

**Figure 5.**
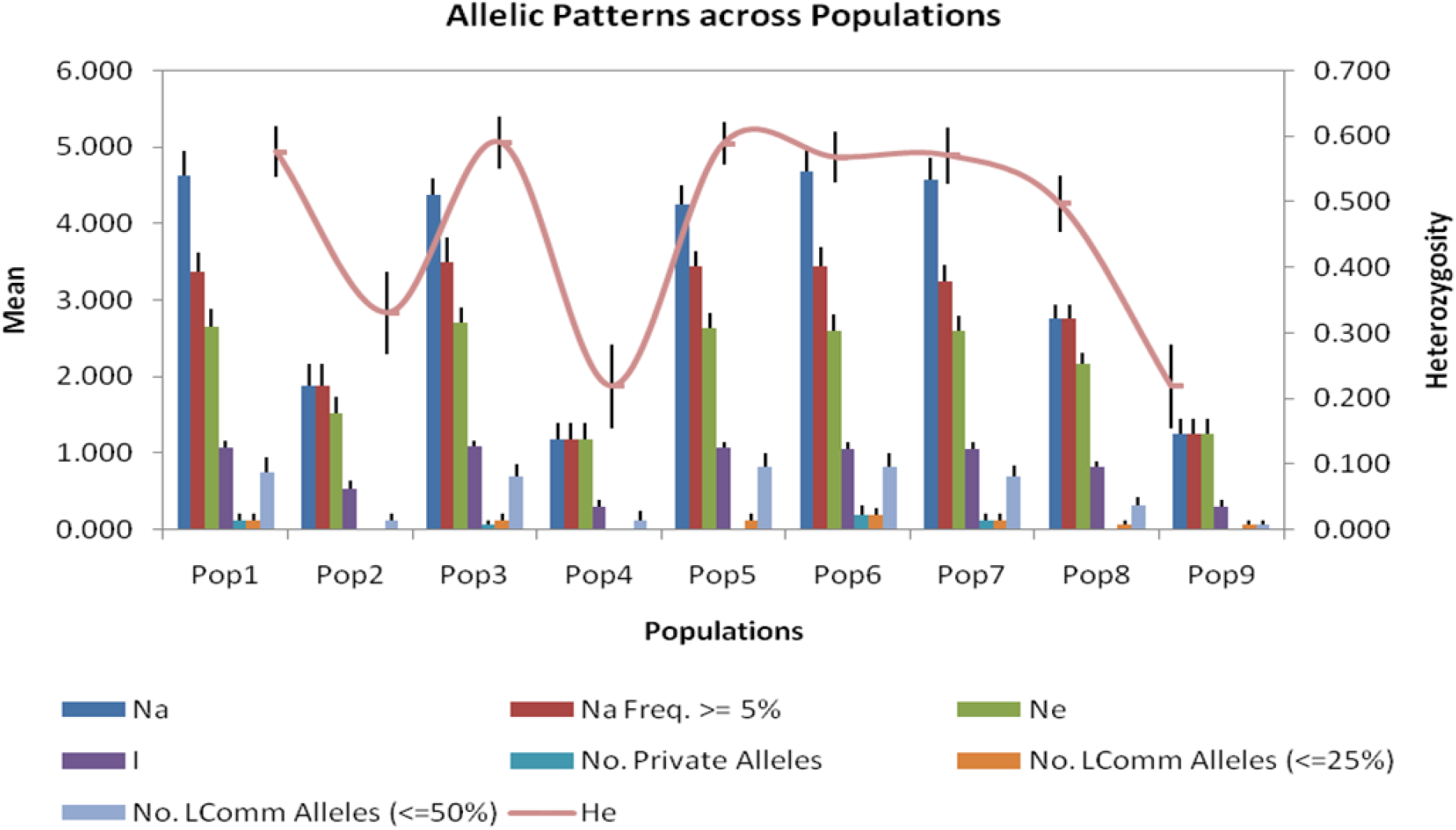
Summary statistics grouped by subpopulations, where *N*_*a*_ represents the number of different alleles; *Na (Freq >= 5%)* the number of Different Alleles with a Frequency ≥ 5%, *N*_*e*_ the number of Effective Alleles, which is equal to 1 / (Σ*p*_*i2*_); I, the Shannon and Weaver Information Index, calculated as Σ(*p*_*i*_ *ln* (*p*_*i*_)); *No. Private Alleles,* the number of Alleles Unique to a Single Population; *No. LComm Alleles (<=25%*), the number of Locally Common Alleles (Freq. >= 5%) found in 25% or fewer populations; *No. LComm Alleles (<=50%)*, the number of Locally Common Alleles (Freq. >= 5%) found in 50% or fewer populations; *H*_*e*_ the Expected Heterozygosity; and *UH*_*e*_ the Unbiased Expected Heterozygosity, estimated as *H*_*e*_(2*N* / (2*N*-1)).

The analysis of allele frequencies identifies a significant number of gaps in the distribution of allele length or number of repeats. It has been reported that populations that have experienced genetic bottlenecks tend to exhibit such less cohesive distributions than stable populations[18].

### Bottleneck Analysis

The microsatellite allele frequency data was tested for departure from mutation-drift equilibrium with the software BOTTLENECK 1.2[19].The results of the three tests of heterozygosity excess (Infinite Allele Model, IAM; Stepwise mutation Model, SMM; and Two-Phase Mutation Model, TPM) are shown in Table 9 and the results of the test for null hypothesis under Sign Test, Standard Difference Test and Wilcoxon Test in Table 10.

**Table 9.** Bottleneck heterozygosity excess test results based on 16 identified loci, where *n* represents the sample size; and *k*_*o*_, the observed number of alleles under the assumption of mutation-drift equilibrium. The IAM, SMM and TPM mutation models simulate the coalescent processes of *n* genes. *H*_*exp*_ is the average heterozygosity and used to compare with the observed value in determining a heterozygosity excess or deficit at each locus. The standardised difference for each locus is estimated based on the inverse product of the Nei gene diversity and standard deviation (SD) of the mutation-drift equilibrium.

| Observed |  |  | IAM |  |  | TPM |  |  |  |  | SMM |  |  |  |  |
| --- | --- | --- | --- | --- | --- | --- | --- | --- | --- | --- | --- | --- | --- | --- | --- |
| locus | *n* | *k*<sub>o</sub> | *H*<sub>e</sub> | *H*<sub>exp</sub> | *SD* | *D*<sub>H</sub>/*SD* | *Prob* | *H*<sub>e</sub> | *H*<sub>exp</sub> | *SD* | *Prob* | *H*<sub>exp</sub> | *SD*<sub>s</sub> | *D*<sub>H</sub>/*SD* | *Prob* |
| VHL20 | 804 | 5 | 0.7 | 0.4 | 0.19 | 1.56 | 0.02 | 0.51 | 0.15 | 1.21 | 0.06 | 0.64 | 0.09 | 0.67 | 0.28 |
| HTG4 | 804 | 4 | 0.43 | 0.33 | 0.2 | 0.52 | 0.37 | 0.42 | 0.17 | 0.11 | 0.47 | 0.55 | 0.12 | -0.96 | 0.16 |
| AHT4 | 804 | 7 | 0.53 | 0.51 | 0.18 | 0.13 | 0.48 | 0.63 | 0.12 | -0.89 | 0.16 | 0.75 | 0.06 | -3.8 | 0.01 |
| HMS7 | 804 | 6 | 0.75 | 0.45 | 0.19 | 1.62 | 0.01 | 0.57 | 0.14 | 1.34 | 0.03 | 0.71 | 0.07 | 0.7 | 0.25 |
| HTG6 | 74 | 4 | 0.68 | 0.46 | 0.17 | 1.37 | 0.05 | 0.53 | 0.14 | 1.1 | 0.1 | 0.6 | 0.1 | 0.8 | 0.21 |
| AHT5 | 804 | 7 | 0.67 | 0.5 | 0.18 | 0.94 | 0.19 | 0.63 | 0.11 | 0.31 | 0.45 | 0.74 | 0.06 | -1.2 | 0.1 |
| HMS6 | 804 | 5 | 0.57 | 0.39 | 0.2 | 0.9 | 0.22 | 0.51 | 0.16 | 0.42 | 0.4 | 0.63 | 0.1 | -0.64 | 0.2 |
| ASB2 | 800 | 7 | 0.57 | 0.5 | 0.18 | 0.43 | 0.41 | 0.63 | 0.12 | -0.43 | 0.26 | 0.74 | 0.06 | -2.73 | 0.02 |
| HTG10 | 804 | 4 | 0.68 | 0.33 | 0.2 | 1.77 | 0.02 | 0.43 | 0.17 | 1.45 | 0.03 | 0.55 | 0.12 | 1.08 | 0.09 |
| HTG7 | 78 | 3 | 0.5 | 0.33 | 0.18 | 0.9 | 0.26 | 0.41 | 0.16 | 0.57 | 0.37 | 0.46 | 0.14 | 0.28 | 0.48 |
| HMS3 | 802 | 6 | 0.2 | 0.45 | 0.19 | -1.36 | 0.14 | 0.58 | 0.13 | -2.89 | 0.02 | 0.7 | 0.08 | -6.29 | 0 |
| HMS2 | 76 | 5 | 0.54 | 0.53 | 0.16 | 0.09 | 0.45 | 0.6 | 0.12 | -0.52 | 0.25 | 0.68 | 0.08 | -1.7 | 0.07 |
| ASB17 | 634 | 9 | 0.72 | 0.59 | 0.15 | 0.84 | 0.21 | 0.71 | 0.09 | 0.08 | 0.45 | 0.8 | 0.05 | -1.83 | 0.05 |
| ASB23 | 742 | 5 | 0.72 | 0.41 | 0.19 | 1.66 | 0.02 | 0.51 | 0.15 | 1.4 | 0.03 | 0.63 | 0.1 | 0.93 | 0.14 |
| LEX3 | 668 | 5 | 0.45 | 0.4 | 0.19 | 0.24 | 0.49 | 0.52 | 0.15 | -0.44 | 0.27 | 0.64 | 0.09 | -2.15 | 0.04 |
| LEX33 | 646 | 8 | 0.76 | 0.56 | 0.17 | 1.14 | 0.08 | 0.68 | 0.1 | 0.79 | 0.21 | 0.77 | 0.05 | -0.34 | 0.29 |

**Table 10.** Tests for null hypothesis under three microsatellite evolution models

| TEST/MODEL | IAM | TPM | SMM |
| --- | --- | --- | --- |
| Sign Test: Number of loci with heterozygosity excess (probability) | 8.93*<br>(0.00120) | 9.40<br>(0.29262) | 9.43<br>(0.06923) |
| Standard differences test: Ti values (probability) | 3.186*<br>(0.00072) | 0.902<br>(0.18357) | 4.294*<br>(0.00001) |
| Wilcoxon Rank Test (probability of heterozygosity excess) | 0.00042* | 0.11560 | 0.97116 |
| *Rejection of null hypothesis (bottleneck) $P < 0.05$ | | | |

Under the Sign Test, the expected number of loci with heterozygosity excess were 8.93 (*p* = 0.00120) under IAM, 9.40 (*p* = .0.29262) under TPM, and 9.43 (*p* = 0.06923) under SMM. This suggests that the null hypothesis is rejected under IAM, but with p> 0.05 would appear to be met under the other two tests. Therefore, only under the IAM is there clear evidence of a recent bottleneck event.

The standard difference test gives T2 probability statistics of 3.186 *(p= 0.00072)* under IAM; 0.902 *(p= 0.18357*) under TPM and −4.294 *(p=0.00001*) under SMM. Probability values of less than 0.05 for both IAM and SMM under these two models suggest a recent bottleneck event.

Under the Wilcoxon rank test the probability values were 0.00042 (IAM); 0.11560 (TPM) and 0.97116 (SMM), thus rejecting the null hypothesis under IAM.

## MODE SHIFT INDICATOR

The Bottleneck software [19] provides an alternative method for detecting potential genetic bottleneck events in the Mode Shift Indicator. Populations that have not experienced a bottleneck will be at or near mutation drift equilibrium and will be expected to have a large proportion of alleles with low frequency (Luikart and Cornuet, 1998). This pattern will show as a normal, L shaped distribution when displayed graphically. Figure 6 shows that the Cleveland Bay data displays a normal L-shaped distribution at low allele size class, but deviates from it in the latter quartiles. This would suggest a population not completely at mutation drift equilibrium, and showing evidence of having experienced a genetic bottleneck in the recent past.

**Figure 6.**
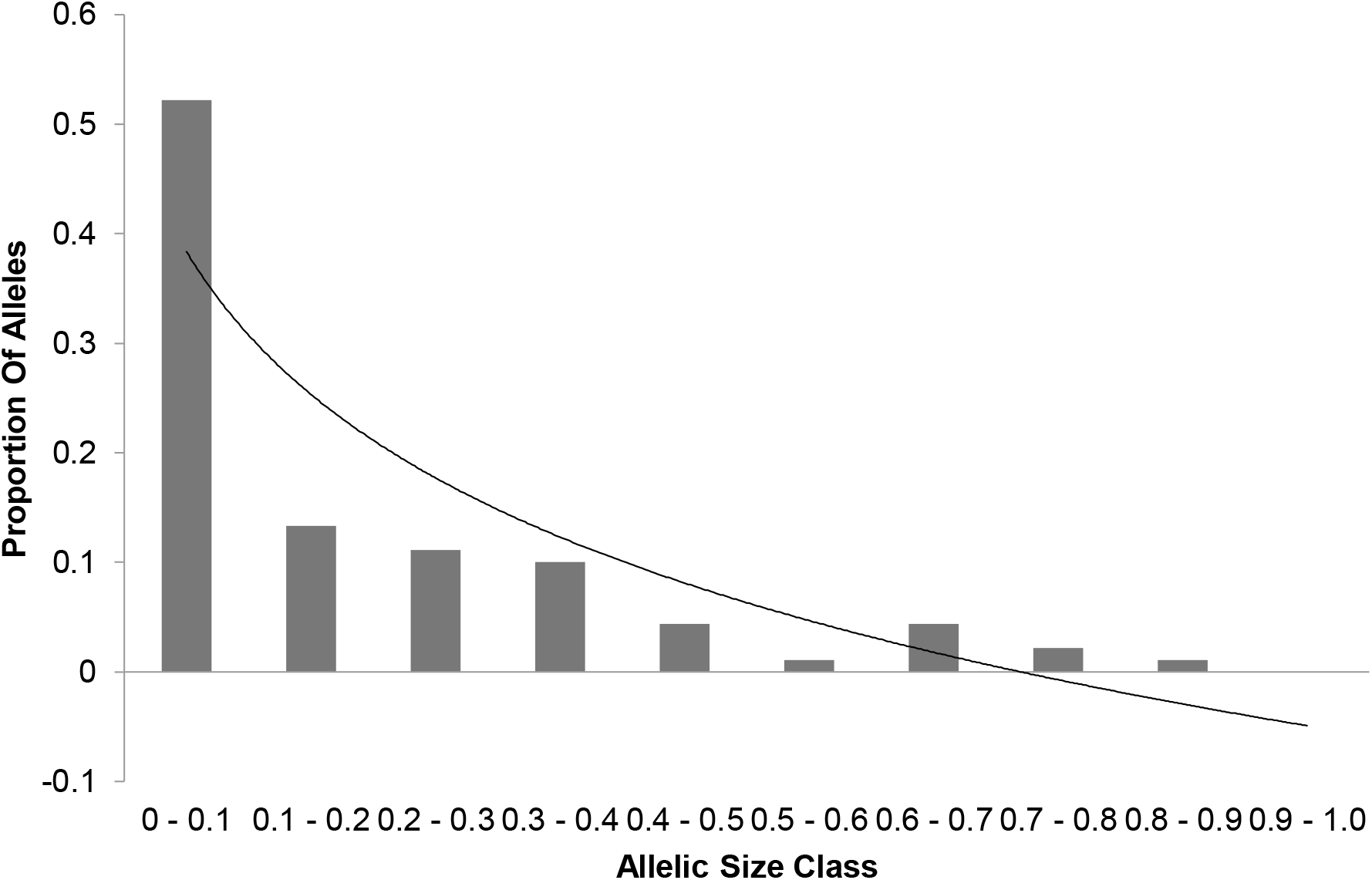
Allele distribution by size class. Trendline describes a natural logarithmic relationship according to *y*=-0.188 *ln*(x) + 0.3836.

As both the data plot and the trend show that at the higher size classes there is some departure from the normal L-shaped distribution; the absolute assumption of accepting the null hypothesis should be treated with caution. Indeed, on initial examination, the results of the analysis with Bottleneck [19] appear far from conclusive. Initial assessment suggests that under the IAM all of the tests provide evidence of a recent bottleneck event. However, under TPM and SMM, the evidence is somewhat contradictory indicates some reservation to assessment of the suggested recent bottleneck. The mutation drift model deviation from normal L-shaped distribution supports the above assumption, however, this conflicting evidence suggests the reduction in population size in the 1950s was perhaps not as significant a bottleneck event as previously reported. When the theory behind the various models is re-examined (Luikart and Cornuet, 1998) it becomes evident that gene diversity excess has only been demonstrated for loci evolving under the Infinite Allele Model. Given that there is very strong evidence to support a recent bottleneck event under this model, which is supported by testing of microsatellite allele frequency data herein, it is likely that the Cleveland Bay horse has indeed experienced a recent genetic bottleneck.

### Population Structure

Wright F Parameters [20] reflecting departure from Hardy-Weinberg equilibrium were calculated from the pedigree analysis for the reference population in terms of *F*_*IS*_ (_-_0.006677), *F*_*ST*_ (0.040230) and *F*_*IT*_ (0.033821). Multilocus estimations of Wright’s F statistics [21] from the microsatellite data showed an across population distribution of the following: *F*_*IS*_ (0.011362), *F*_*IT*_ (0.029308), and *F*_*ST*_ (0.018153).

Distance matrices [22] were constructed from both pedigree and molecular analysis, and phlogenetic trees were constructed using TRex [23] showing the relative positions of each female ancestry line (Figures 7 and 8).

**Figure 7.**
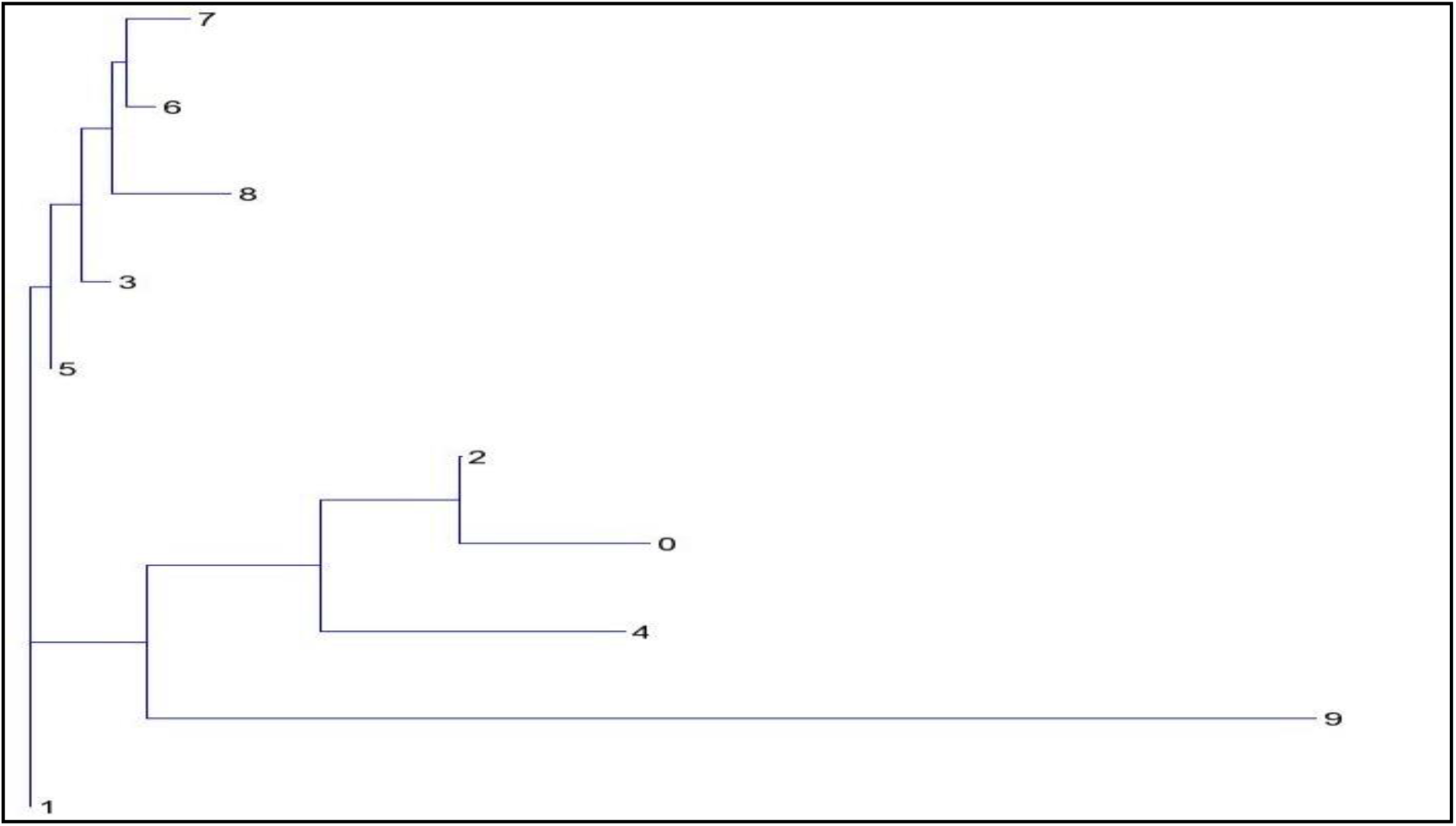
Neighbour Joining Tree showing relative genetic distance between subgroups from analysis of pedigree data assigned by female ancestry line.

**Figure 8.**
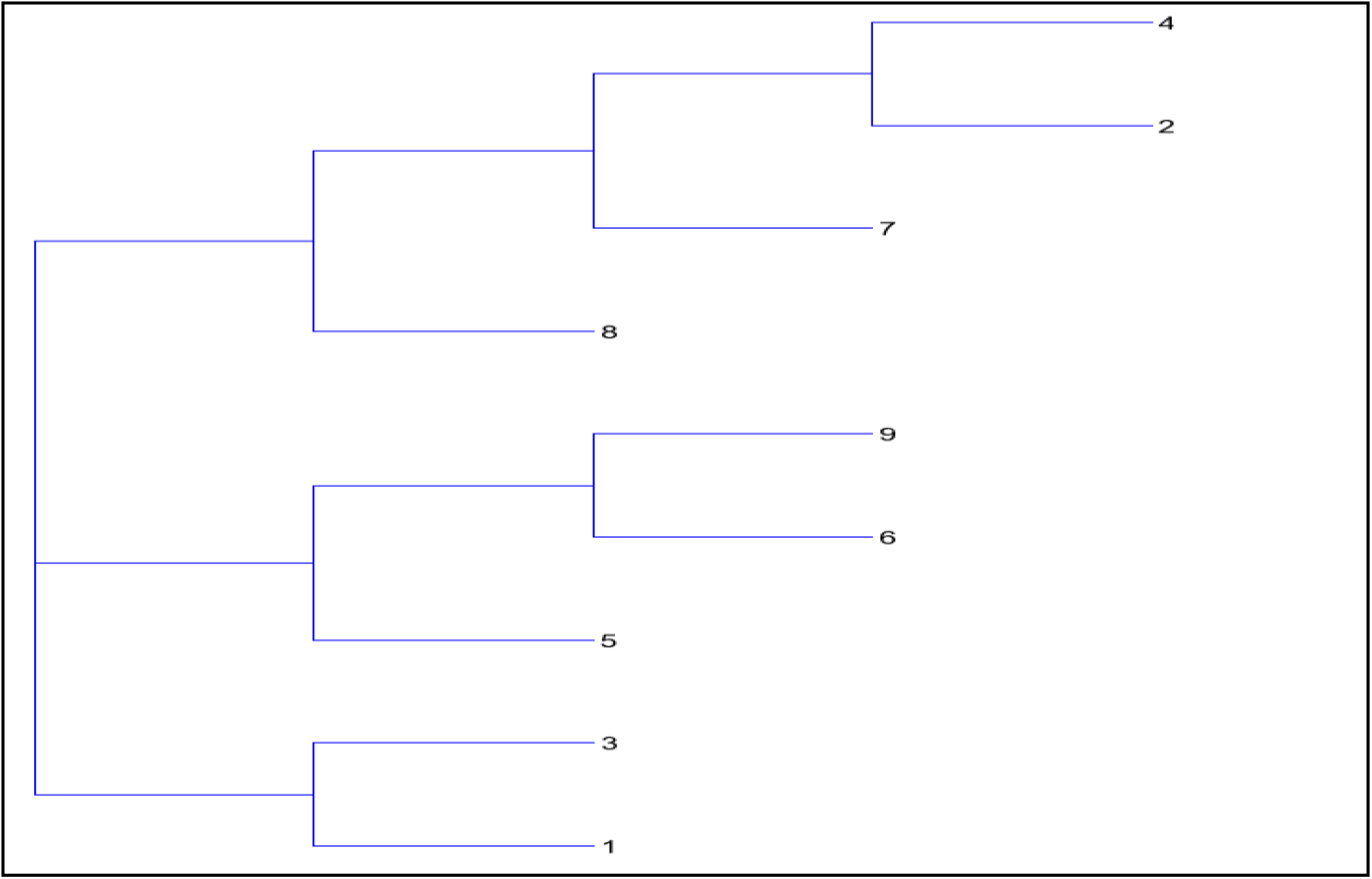
Neighbour Joining Tree from Microsatellite analysis showing distances between subpopulations by maternal ancestry line.

Both the pedigree distance analysis (Figure 7) and the molecular analysis (Figure 8) are suggestive of a population structure rooted on three sub-divisions, or clades. However, neither analysis provides conclusive evidence of the causes or nature of this division. In addition to the pairwise distance matrices constructed assuming 9 subgroups within the population, GENALEX 6.4 [24] was also used to construct the much larger matrix of Nei distance between individuals [22]. This matrix in Phylip format was imported into the cluster drawing programme SplitsTree4 [25] to construct a Neighbour-Net diagram (Figure 9).

**Figure 9.**
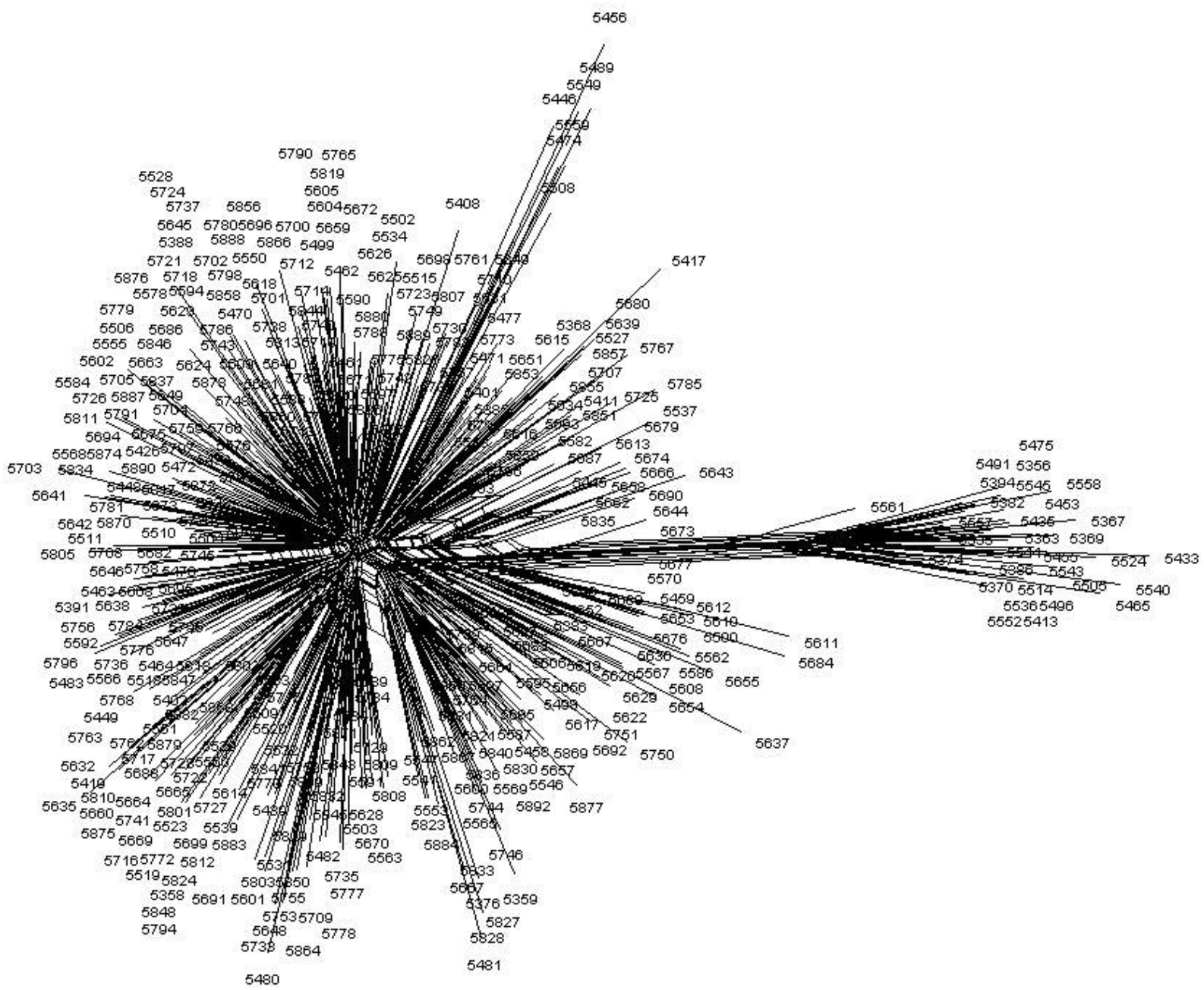
Neighbour-Net diagram of Nei genetic distance between individuals.

Examination of this net immediately suggests that the structure of the reference population could be explained by two broad groups or clades as shown in Figure 10. However, an alternative model with three clades, shown in Figure 11, is also possible.

**Figure 10.**
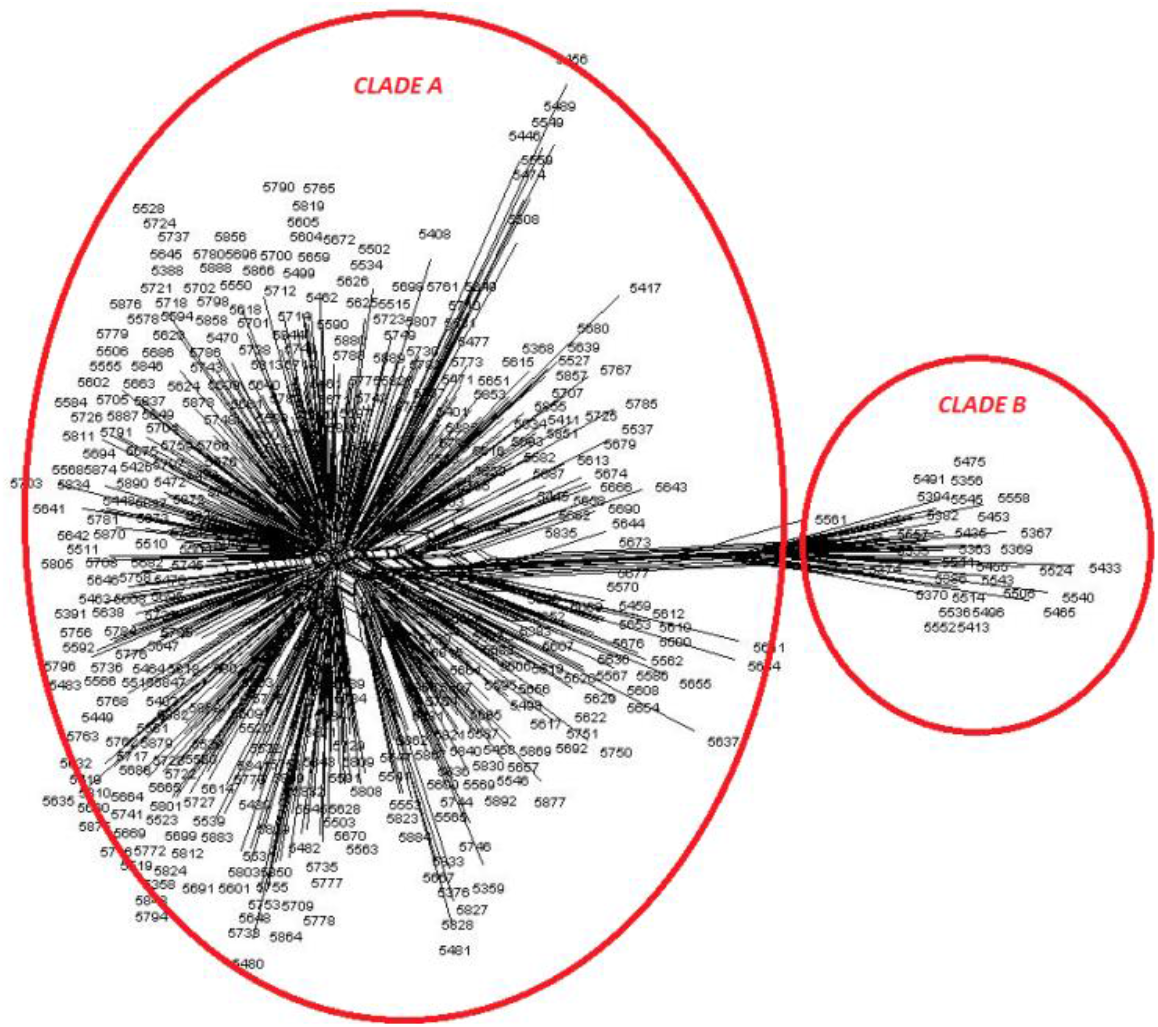
Neighbour-Netdiagram of Nei genetic distance between individuals showing two clade model of structure.

**Figure 11.**
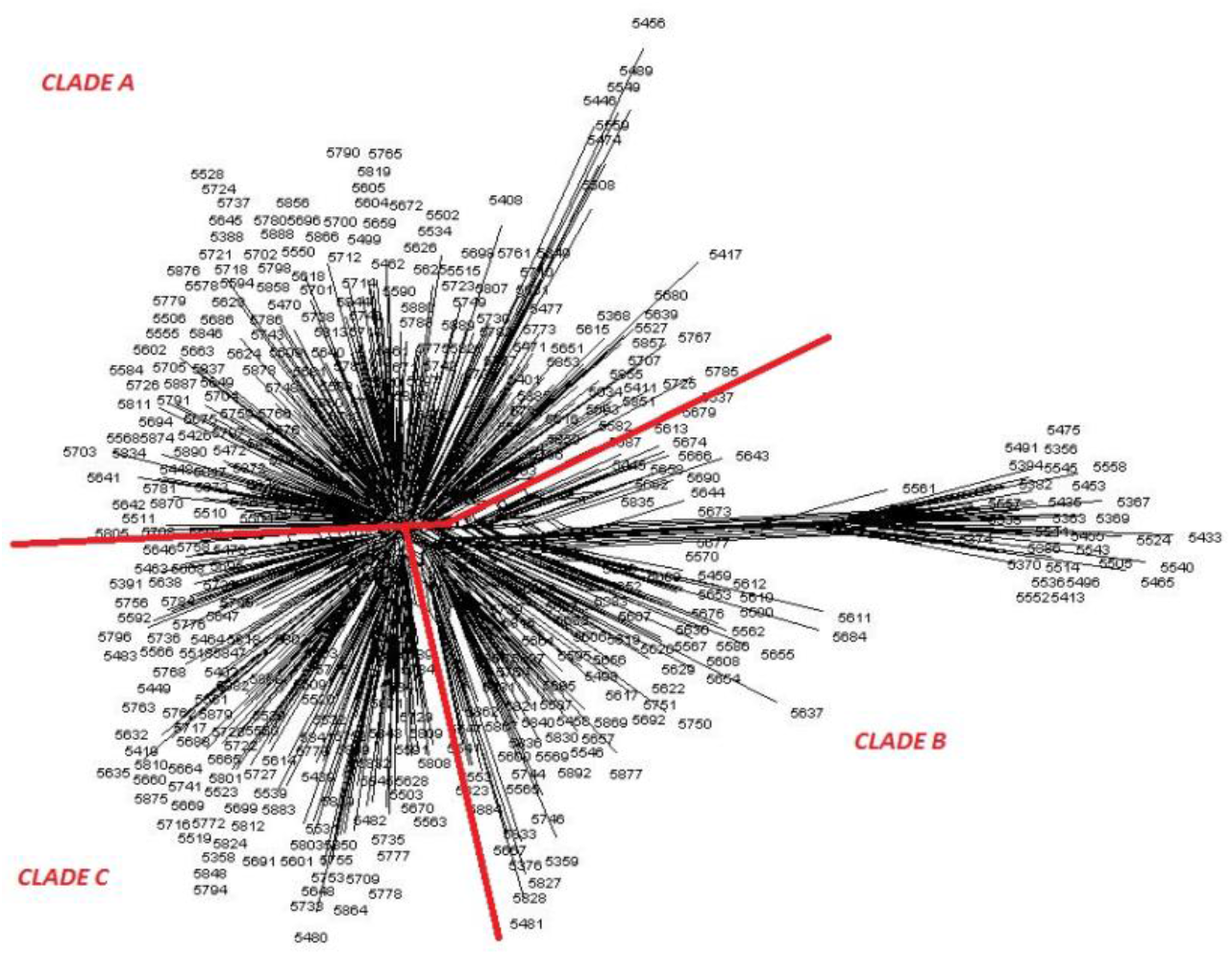
Neighbour-Netdiagram of Nei genetic distance between individuals showing three clade model of structure

Principal co-ordinate analysis *via* covariance matrix was conducted using Genalex 6.5 [24], with sub-populations assigned by both modern female and modern male ancestry lines, in order to examine alternative possible structuring of the reference population. Figure 12 presents the PCoA with subpopulations assigned by female ancestry and Figure 13 by male ancestry.

**Figure 12.**
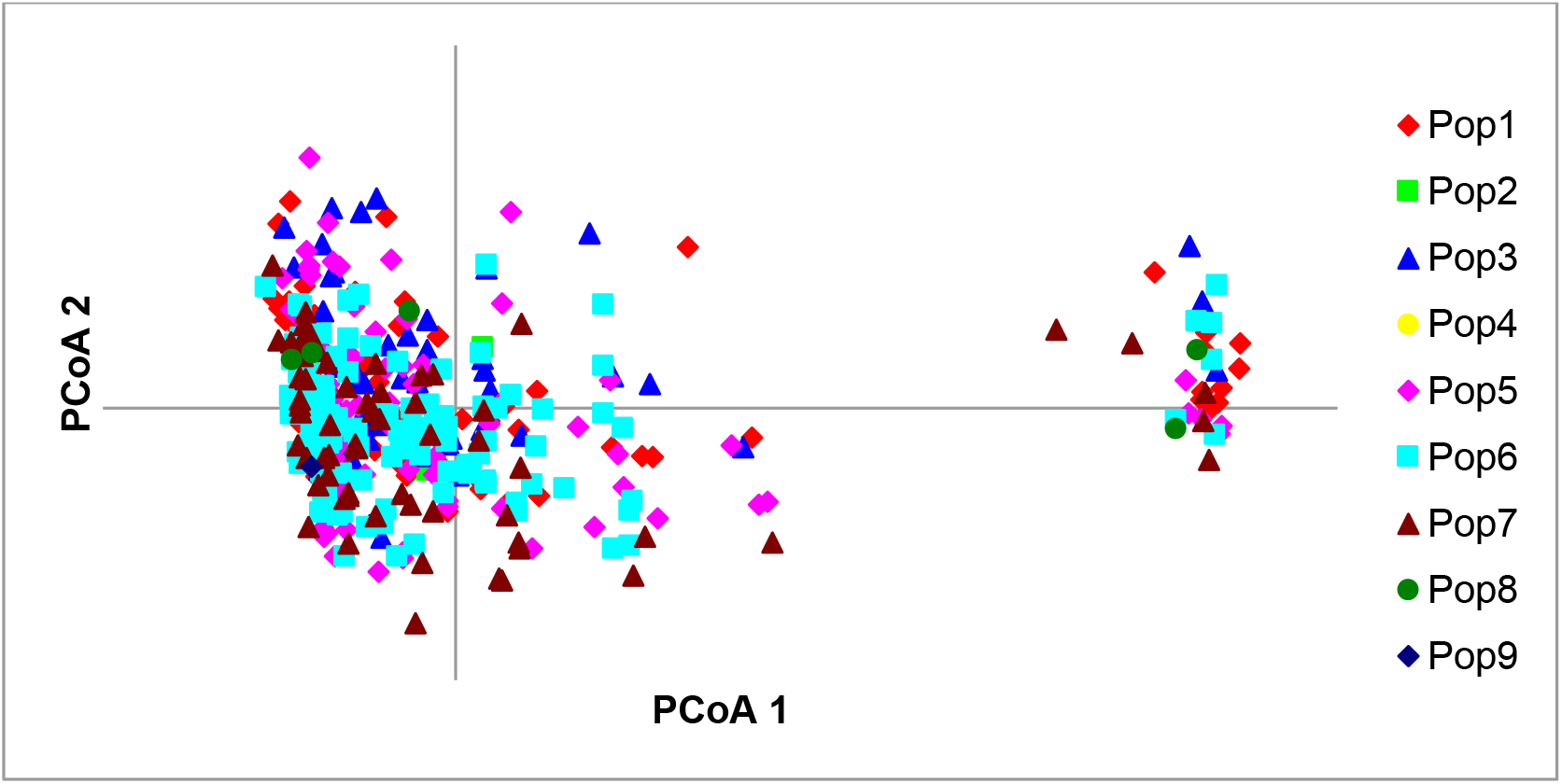
Principal Coordinate Analysis (PCoA) with subpopulations assigned by female ancestry across the two principal components (PCoA1, PCoA2).

**Figure 13.**
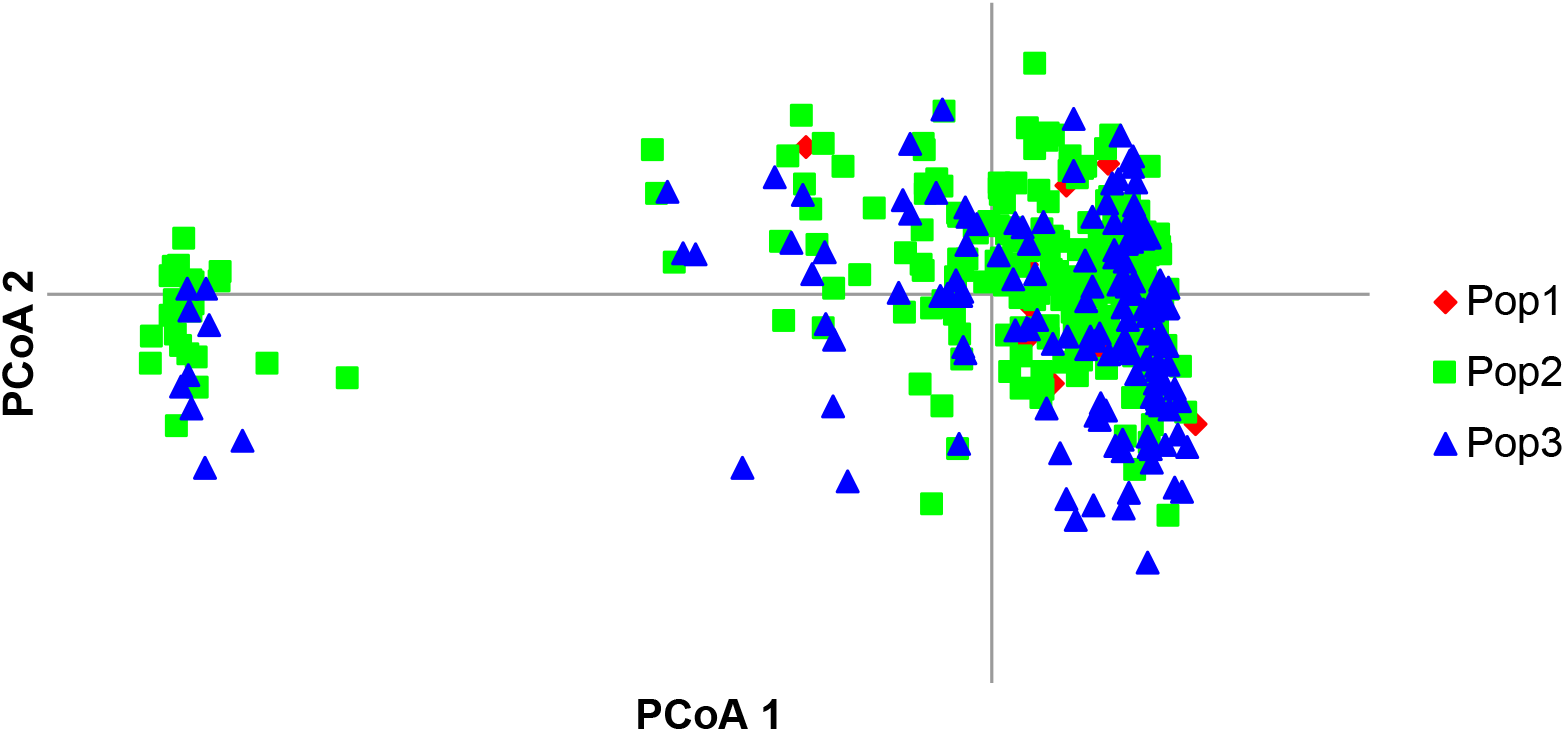
PCoA with subpopulations assigned by male ancestry.

The PCoA analysis shows both male and female sub-populations distributed widely across principal axes, with little suggestion of structuring by sex group being the driving process of population sub-division in the microsatellite data. Variational Bayesian analysis of the microsatellite dataset, using the programme STRUCTURE [26] was carried out, in order to further investigate breed structure. 104 runs of the analysis were carried out for potential populations, *K*, numbering 2 to 25. The best fit of *K* appears at *K = 3*. Figure 14 provides a visual representation of this analysis for *K = 2* to *K = 4*. There is a substantial increase in background noise in the display at *K = 4*, indicative that the number of clusters or sub-populations is below this level.

**Figure 14.**
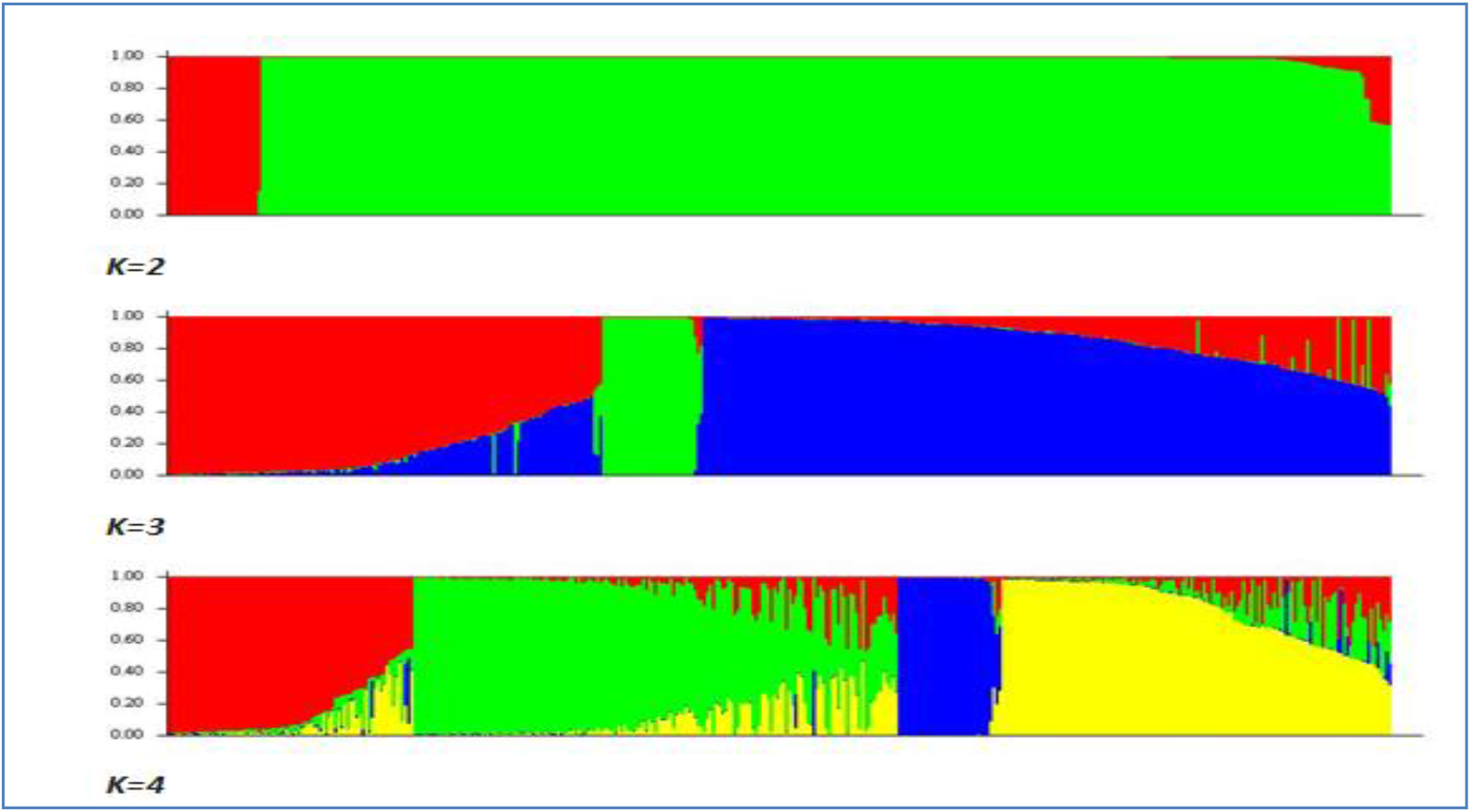
STRUCTURE analysis of population numbers *K=2* to *K=4*. Each colour is a representation of a population, with individuals shown as vertical lines, which are split into coloured segments; the lengths of these describe the admixture proportions from *K* populations.

Further analysis of the population structure was conducted using the programme BAPS[27]. 17 clusters within the microsatellite dataset were identified, with a highly significant probability of 0.99998.

## Conclusions

The results presented herein highlight the significant losses of founder representation that have occurred in the Cleveland Bay Horse population across the past century. Approximately 91% of the stallion and 48% of the dam lines are lost in the reference population. The unbalanced representation of the founders is illustrated by the effective number of founder animals (*f*_*e*_) and the effective number of ancestors (*f*_*a*_). The parameter *f*_*e*_ constitutes over a third of the equivalent number of founder animals for the reference population, whilst the ratio *f*_*a*_/*f*_*e*_ is 22.5%. This ratio is substantially lower than that reported in other horse breeds such as 41.7% in the Andalusian [28] or 54.4% in the Lipizzan [29]. Additionally, this is lower than the figure of 38.2% reported for the endangered Catalonian donkey [30].

The average inbreeding computed for the Cleveland Bay Horse at 20.64% in the reference population is substantially higher than most of the values reported in the literature[28], with typical values ranging from 6.5% to 12.5%. Although most of these inbreeding values have been computed in breeds with deep pedigrees such as Andalusian, Lipizzan or Thoroughbred there are significant differences in population sizes, and the accumulation of inbreeding in populations of restricted size will occur at a greater rate.

The smaller the number of individuals in a randomly mating breed the greater will be the accumulation of inbreeding due to the restricted choice of mates. The Cleveland Bay horse is therefore predisposed to inbreeding and associated loss of genetic variation. In the reference population of 402 individuals the Effective Population Size (*N*_*e*_) computed *via* individual increase in inbreeding was 27.84. *N*_*e*_ computed *via* regression on equivalent generations was 26.29. Inbreeding and genetic loss under random mating will occur at ½ *N*_*e*_ per generation. In the reference population, where Mean *N*_*e*_ is 32.32 under random mating, inbreeding can be expected to accumulate at 1.5% per generation.

This is reflected by the genealogical *F*_*IS*_ values. This parameter characterises the mating policy derived from the departure from random mating as a deviation from Hardy-Weinberg equilibrium. Positive *F*_*IS*_ values indicate that the average *F* value within a population exceeds the between-individuals coancestry, thus suggesting that matings between relatives have taken place [30]. Moreover, the average AR values computed for nine complete generations, as shown in Table 3, are roughly equivalent the value of *F*. In an ideal scenario with random matings and no population subdivision, AR would be approximately twice the *F* value of the next generation [30].

Molecular information obtained in this study using microsatellite analysis suggests that genetic diversity within the breed is more restricted than has been reported in many other horse breeds.

Populations that have experienced a recent reduction in their *N*_*e*_ exhibit a correlative reduction of the allele numbers (*k*) and gene diversity (*H*_*e*_) at polymorphic loci. However, the allele numbers reduce faster than the genetic diversity. Thus, in a recently bottlenecked population, the observed gene diversity is higher than the expected equilibrium gene diversity (*H*_*e*_) which is computed from the observed number of alleles, *k*, under the assumption of a constant-size or equilibrium population [43].The existence of a population bottleneck in the mid twentieth century, when the number of breeding age Cleveland Bay stallions was reduced to four, has previously been reported [13]. There is clear genetic evidence of this event shown in the excess of observed heterozygosity across subpopulations, with the exception of ancestry line nine. The latter is of more recent origin having evolved from a grading up scheme in the latter half of the twentieth century. In all other subgroups, the excess is positive ranging from 2.12% in Line 5 to 19.6% in Line 4. However, this investigation has revealed that lines two, four, and eight are in fact not polymorphic. The observed heterozygosity excess amongst the five polymorphic lines peaks in line one at 6.1%.

Microsatellite multilocus estimations of Wright’s *F* statistics [21] showed an across population *F*_*IS*_; *F*_*IT*_ and *F*_*ST*_ of 0.01758, 0.02490, and 0.00745, respectively. This departure from random mating will have been influenced by a number of factors common to restricted populations of domesticated equines. These include: selection by breeders for particular lines of descent; natural differences in fertility between individuals; a restricted number of male animals leaving significantly more offspring than females (disproportionate male founding) and geographic distribution of animals and breeders leading to logistical difficulties in some matings. The reduced number of alleles and fixation at certain loci in female ancestry lines is evidence of loss of founder representation from these lines. This lower heterozygosity is also indicative of the typical practice of the larger studs, where breeding tends to be carried out in pasture by free live cover, with the use of only one stallion per year, per herd and where the same stallion may be retained for several breeding years. This strategy is compounded by breeders with only a small number of breeding females sending their animals to these groups or to be covered in hand by the same stallion.

This strategy has different implications for the genetic diversity of the Cleveland Bay Horse compared that of mares travelling to stud to be covered in hand by a greater range of stallions that do not have their own herds of mares [42]. as well as through trade or exchange, which will change geographic location albeit on an irregular basis. Although his latter practice has clear benefits in conservation programmes, there is the danger of inappropriate matings supplanting the more common and less frequent alleles. Whilst such matings increase the frequency of the rarer alleles, they simultaneously increase the frequency of those more common [44], highlighting the need for in-depth understanding of the genetic diversity of any rare breed, and for an effective management plan for conservation maintenance.

There has been considerable debate about the most effective methods of conserving and managing endangered populations [42]. Before the advent of mitochondrial and microsatellite DNA analysis, the accepted strategy involved minimizing inbreeding, whilst managing mean Kinship/average relatedness [14]. Moreover, the use of molecular methods has been proposed [45,46]. Where pedigree data is robust and complete over a significant number of generations, it appears that genealogical data remains the preferred method by which to manage founder contributions, inbreeding and kinship/relatedness. Indeed Lacy has highlighted the problems caused in conservation programmes based on private or rare alleles [44].

Variational Bayesian analysis of within-population structure using microsatellite data shows significant evidence for three main clades. Although this study has been based on the use of pedigree and microsatellite marker data for the Cleveland Bay horse there is now firm evidence of the value of mitochondrial DNA for such investigations and an increasing number of investigations consider the origins and relatedness of modern equines (Table 11). The Cleveland Bay horse has been reported to belong to haplotype C [34] which is common amongst older northern European breeds such as the Exmoor, Icelandic, Fjord, Connemara and Scottish Highland. This correlates with the assertion that in the matriline the Cleveland Bay has evolved from the Chapman; an ancient Northern European breed (Dent, 1978). The comparative studies have been based on five Cleveland Bay mtDNA sequences deposited in GeneBank by Cothran and Frankham within which there are three haplotypes. There is scope for further sampling of all of the existing matrilines to determine the number of haplotypes present in the reference population the level of correlation with the three Clades identified herein.

**Table 11.** Genetic variability from microsatellite DNA loci for Cleveland Bay and other domestic horse breeds. *H*_*e*_ denotes the expected heterozygosity, whilst *H*_*o*_ represents the observed heterozygosity, and MNA the mean number of alleles per locus.

| Breed | He | Ho | MNA | Source |
| --- | --- | --- | --- | --- |
| Cleveland Bay | 0.173- | 0.052-0.716 | 6.19 |  |
| Suffolk Punch | 0.724 | 0.679 | 6.42 | [31] |
| Dales Pony | 0.654 | 0.715 | 5.58 | [32] |
| Exmoor Pony | 0.609 | 0.601 | 5.25 | [33] |
| Fell Pony | 0.731 | 0.782 | 6.42 | [32]oor |
| Irish Draught | 0.772 | 0.766 | 7.08 | [34] |
| Shetland | 0.661 | 0.642 | 5.3 | [35] |
| Thoroughbred | 0.646–<br>0.732 | 0.628–<br>0.671 | 4.7–7.5 | [2] |
| Thoroughbred | 0.695 | 0.674 | 6.25 | [36] |
| Arabian | 0.690 | 0.624 | 6.58 | [37] |
| Lipizzan | 0.675 | 0.663 | 7.1 | [38] |
| Friesian | 0.466 | 0.454 | 4.5 | [39] |
| Spanish Celtic<br>Horses | 0.677–<br>0.770 | 0.694–<br>0.765 | 5.2–7.8 | [40] |
| Portuguese | 0.751 | 0.732 | 4.5 | [39] |
| Lithuanian | 0.442–<br>0.770 | 0.452–<br>0.785 | 2.0–4.7 | [41] |
| Sorraia Horse | 0.093–<br>0.736 | 0.088–<br>0.705 | 3.3 | [42] |

We have reported an in-depth genetic analysis of the Cleveland Bay Horse, using both pedigree and microsatellite data. It reveals substantial loss of genetic diversity and high levels or relatedness and inbreeding. The results of this study highlight the importance of the Cleveland Bay Horse community implementing an effective and sustainable breed management plan, such as management of Mean Kinship and Inbreeding Coefficients.

## Materials and Methods

### Pedigree data

Summary data from the CBHS stud books volumes one to thirty eight was published in the Society’s Centenary studbook [7]. Names and studbook numbers of all registered horses together with date of birth, sire and dam were listed and this information was digitised in Filemaker™ (Filemaker Inc.), to construct an electronic pedigree data base for the breed, stored in Filemaker format. Registrations post-1985 have been added to the database on an annual basis up to and including for this study, Volume 38 of the studbook.

The Cleveland Bay Horse Society provided access to a total of 535 microsatellite parentage testing reports. These had been obtained by commercial analysis of hair follicle samples taken from individual animals for registration verification. Samples were tested for a panel of 16 microsatellite markers approved by the International Society for Animal Genetics (ISAG) equine genetics group, by the Animal Health Trust (Newmarket, UK.). Close examination of stud book records, recent Breed Society census records and the microsatellite dataset enabled the identification of a reference population of 402 animals, registered in the 10-year period 1997 to 2006.

### Pedigree Completeness

Data correction routines within the programmes Genes [47]and Eva [48] were used to identify pedigree errors and correct infinite loops. Calculation of Pedigree Completeness was made using PopRep [17]Using Equations 1 and 2 to compute pedigree completeness index [49](I_d_):

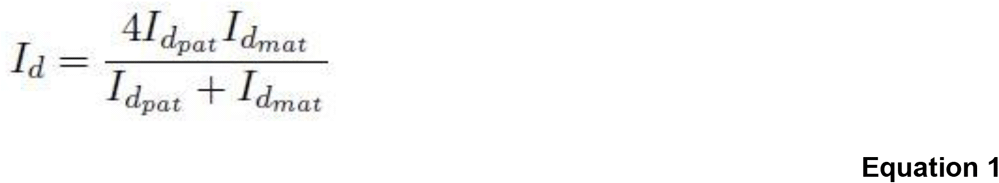

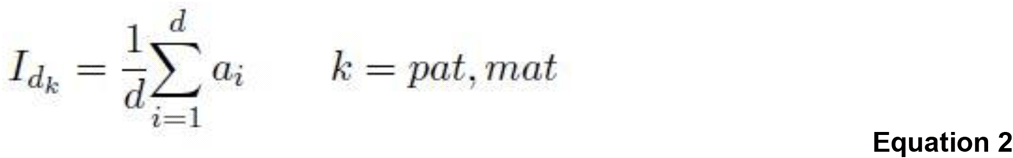

Where *k* represents the paternal (pat) or maternal (mat) line of an individual, and *a*_*i*_ is the proportion of known ancestors in generation *i*; *d* is the number of generations measured when calculating the pedigree completeness. Values for pedigree completeness will range from 0 to 1. Where all of the ancestors of an individual are known to some specified generation (*d*) then *I*_*d*_=1. However, where one of the parent animals is unknown, *I*_*d*_ = 0[17].

### Generation Interval

Generation Interval is defined as the average age of the parent animals at the birth of selected offspring with offspring subsequently producing at least one progeny [50]. The generation interval was calculated for each of the four possible lines of descent: sire to son; sire to daughter; dam to son and dam to daughter. The results were averaged for each year group using PopRep [17].

### Founder and Ancestor Representation

Stallion and dam lines, defined respectively as: *unbroken descent through male or female animals only from an ancestor to a descendant* [3] were identified and detailed founder and ancestor analysis was performed using Endog 4.6 [51] to initially determine Number of Founders.

We make the assumption that all animals with two unknown parents are regarded as founders in this analysis. In addition, if an animal has one known and one unknown parent, the unknown parent is regarded as a founder. The total number of founders contains limited information on the genetic basis for the population. Firstly, founders are assumed to be unrelated, as their parentage is unknown. However, this is most likely not the case in practice. Secondly, some founders have been used more intensely and therefore contribute more, in terms of genetic resource, to the current population than other founders. The effective number of founders, *f*_*e*_, has been designed to correct for this second shortcoming.

The Effective Number of Founders (*f*_*e*_) [52] is defined as the number of equally contributing founders that would be expected to produce the same genetic diversity as in the population under study. This is computed as:

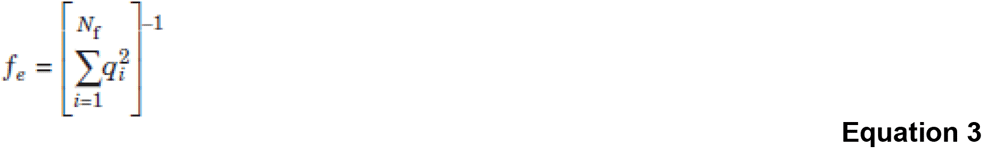

Where *q*_*k*_ is the probability of gene origin of the *k*_th_ founder and *N*_*f*_ the real number of founders. In a scenario where every founder makes an equal contribution, the effective number of founders will equal the actual number of founders.

It is more common for founders to contribute unequally, leading to *f*_*e*_ < *N*_*f*_. The genetic contributions will converge following 5 to 7 generations[53]. Once this convergence occurs, employing *f*_*e*_ as a measure of genetic contribution, will have limited usefulness as will remain constant irrespective of later changes in the population. Pedigrees of more than 7 generations can be characterized with a high effective number of founders even after a severe, recent bottleneck [19]. Whilst the effective number of founders is not an absolute measure of genetic diversity, it forms a basis for comparison of the effective population size (*N*_*e*_) and the effective number of ancestors). In a population with minimum inbreeding, *f*_*e*_ would be expected to be approximately equal to ½*N*_*e*_ [53]. Where *f*_*e*_ diverges from this, there is compelling evidence that the breeding structure has been changed since the founder generation [54].

The Effective Number of Founder Genomes (*f*_*g*_) was proposed by Lacy (1989) to account for unequal founder contributions, random loss of alleles caused by genetic drift and for bottleneck events. It is computed by the equation:

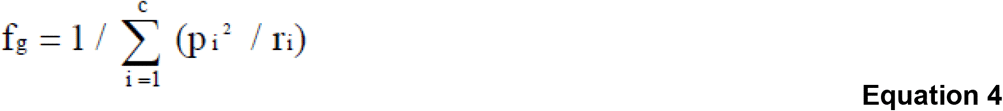

Where *p*_*i*_ is the expected proportional genetic contribution of a founder *i*; *r*_*i*_ is the expected proportion of alleles from founder *i* which remain in the current population, and *c* is the total number of contributing founders[52]. This gives an indication of the number of equally contributing founders with no loss of founder alleles, that would produce the same degree of diversity as found in a reference population[55]. The *f*_*g*_ will be smaller than both *f*_*e*_ and the effective number of ancestors (*f*_*a*_), even under minimum inbreeding pressure, and approximately equal to ½*N*_*e*_. The scale of these differences is indicative of the degree of random loss of alleles. Alleles will be lost with every generation of a pedigree and thus *f*_*g*_ will decrease as the depth of pedigree increases [54].

The Effective Number of Ancestors (*f_a_*) supplements *f*_*e*_ and is calculated from the genetic contributions of ancestors with the largest marginal genetic contributions themselves [16]. Whilst genetic contributions of founders are independent and sum to unity, this is not the case for genetic contributions of ancestors. Indeed, the dam of a highly used sire has >50% contribution of her son, as the same genes are represented in both generations. Boichard et al. (1997) therefore introduced the marginal contribution to the pedigree genetic resource. The ancestors contributing most to the reference population are considered individually in a recursive process. For each round of the recursion, the ancestor with the highest contribution is chosen, and the contributions of all others are calculated conditionally on the contribution of the chosen ancestor. The marginal contribution is the genetic contribution from an individual after correcting for contributions of other ancestors already considered in the recursive process. The sum of marginal contributions of all ancestors will be equal to unity. Ancestors with a large marginal contribution to the reference population will correlate with individuals having genes passed through many descendants [54].

Assessment of the *f*_*a*_ helps to account for the losses of genetic variability produced by the unbalanced use of individuals in terms of reproduction within breeding programmes. This is conventional in domestic equines, whilst also accounting for bottlenecks in the pedigree.

The parameter *f*_*a*_ is computed as

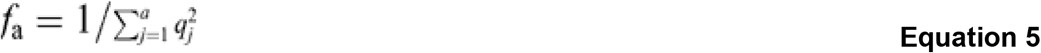

where *q*_*j*_ is the marginal contribution of an ancestor *j*.

### Inbreeding Analysis

Inbreeding coefficients for each individual animal were calculated using ENDOG[51].

The Increase in Inbreeding (Δ*F*), is calculated for each generation using ENDOG 4.6[51], by means of Equation 6.

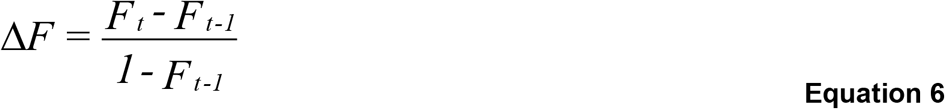

where *F*_*t*_ and *F*_*t-1*_ are the average inbreeding of offspring and their parents, respectively[50].

The Average Relatedness Coefficient (*AR*) [30] describes the probability that a randomly chosen allele from the whole population in the pedigree belongs to the animal under study. This parameter was calculated using ENDOG 4.6 [51].The Additive Relationship Coefficient (R_yz_), is estimated for two animals through calculating the hypothetical coefficient of inbreeding of an animal produced by mating the two individuals, irrespective of the sex of these assumed parents. The additive relationship between the two animals is then calculated as twice the coefficient of inbreeding of the hypothetical offspring. R_yz_ = 2 *F*_*x*_, where *F*_*x*_ is the coefficient of inbreeding of the hypothetical offspring of individual Y and individual Z. This additive relationship has a minimum value of zero and a maximum value of two. The Additive Relationship is twice the value of the coefficient of kinship. The kinship of any two individuals is identical to the inbreeding coefficient of their progeny if they were mated. It is the probability that alleles drawn randomly from gametes of each of the two individuals are identical by descent.

### Effective Population Size

The Effective Population Size from the rate of inbreeding is computed using the classic equation

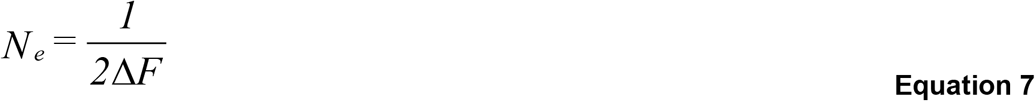

Where the rate of inbreeding per generation is calculated using Equation 6.

The Effective Population Size from the number of parents is computed as

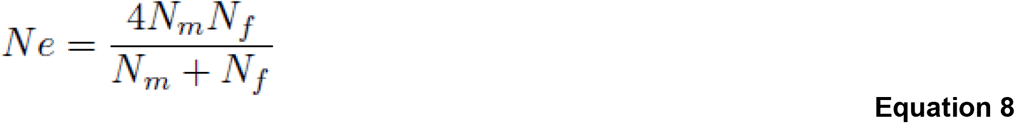

Where *N*_*m*_ and *N*_*f*_ are the number of male and female parents, respectively [50]. This method assumes that the ratio of breeding males to breeding females is 1:1, and that all individuals have an equal opportunity to contribute their genetic material to the next generation. This is seldom the case in managed livestock populations and there is a tendency for this method to overestimate *N*_*e*_ [17].

### Microsatellites

Total DNA was isolated at the Animal Health Trust’s laboratories, from hair follicle samples following standard commercial procedures and as previously described [37]. A set of 16 microsatellites (ASB17 VHL20 HTG10 HTG4 AHT5 AHT4 HMS3 HMS6 HMS7 ASB23 LEX3 LEX33 ASB2 HTG6 HTG7 HMS2) were analysed in all the sampled individuals. The GENETIX program was used to carry out factorial correspondence analyses and associated calculations [56].

The Average Number of Alleles per Locus (*A*), corrected in order to account for sample size using Hurlbert’s rarefaction method (1971) can be shown as:

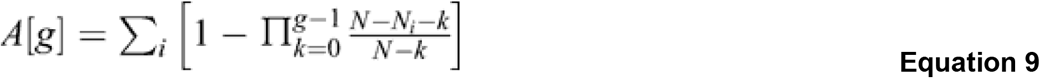

where *g* is the specified sampled size for a collection containing *N* individuals, numbering *N*_*i*_ in the *i*_th_ species.

Nei’s minimum distance (*D*_*m*_) and Nei’s standard distance (*D*_*s*_[57] are computed according to Equations 10 and 11, respectively.

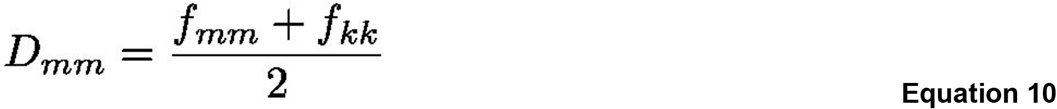

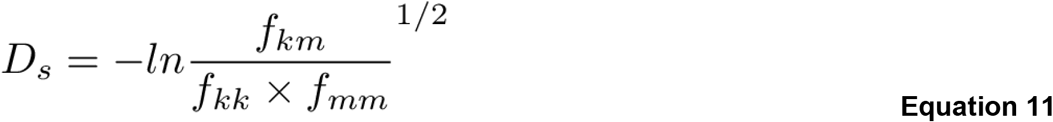

where *f*_*kk*_ and *f*_*mm*_ are the average coancestry between individuals belonging to population *k* or *m*, and *f*_*km*_ is the average coancestry between individuals belonging to populations *k* and *m*.

### Population Structure

*F* (fixation) statistics extend the study of inbreeding coefficients in the case of sub-divided populations [58]. The *F*_*IT*_ refers to the inbreeding of individuals in the total population. Conversely, *F*_IS_ describes the inbreeding of individuals within sub-populations. *F*_*ST*_ is not strictly a fixation index as it represents the correlation between two gametes taken at random in two sub-populations from the total population. It measures the degree of genetic differentiation of the sub-populations. The three indices are computed as in Equations 12, 13, and 14, respectively

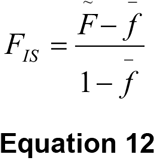

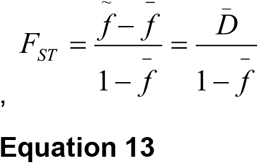

and

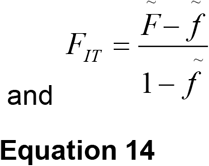

where *f* and *F* are, respectively, the mean coancestry and the inbreeding coefficient for the entire metapopulation, and, the average coancestry for the subpopulation, so that (1 - *F*_*IT*_) = (1 - *F*_*IS*_)(1 - *F*_*ST*_) [59]‥

ENDOG [51] was used to calculate *F* statistics and Nei’s minimum distance[57]), *D,* the genetic distance between subpopulations *i* and *j* which is given by Equation 15

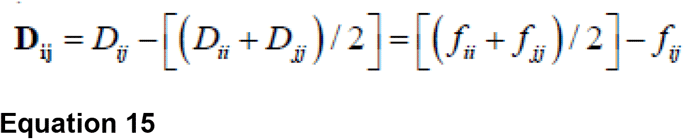

The programme TREX [23,60] was used to construct phylogenetic trees to illustrate the structure from the distance matrix data.

Bayesian model-based clustering was conducted using the programme STRUCTURE v2.1 [26], to assign individuals to homogeneous clusters or populations *K*, from a user defined range. An admixture model was adopted, with a burn in of 104 and 104 iterations of each value of *K* from 2 to 25.

## Acknowledgments

The authors thank the Breed Committee of the Cleveland Bay Horse Society for access to its microsatellite parentage testing records. AD was supported by an MPhil/PhD grant from the Department of Biological Sciences, University of Lincoln, and by the RBST / Marsh Christian Trust Award for Conservation of Genetic Biodiversity 2008.

## References

1. Petersen JL, Mickelson JR, Cleary KD, McCue ME. The american quarter horse: Population structure and relationship to the thoroughbred. Journal of Heredity. 2014. doi:10.1093/jhered/est079

2. Cunningham EP. Molecular methods and equine genetic diversity. Conservation Genetics of Endangered Horse Breeds. 2005.

3. Cunningham EP, Dooley JJ, Splan RK, Bradley DG. Microsatellite diversity, pedigree relatedness and the contributions of founder lineages to thoroughbred horses. Anim Genet. 2001. doi:10.1046/j.1365-2052.2001.00785.x

4. Boyd MM. A plea for a more extended use of the system of live-stock registration. J Hered. 1907. doi:10.1093/jhered/os-3.1.255

5. Pritchard JK, Stephens M, Donnelly P. Inference of population structure using multilocus genotype data. Genetics. 2000.

6. Khanshour AM, Hempsey EK, Juras R, Cothran EG. Genetic characterization of Cleveland Bay Horse Breed. Diversity. 2019. doi:10.3390/d11100174

7. Emmerson S. Cleveland Bay Horse Society Centenary Studbook. Clevel Bay Horse Soc. 1984.

8. Dent AA. Cleveland Bay Horses. JA Allen & Company, Limited; 1978.

9. Fairfax-Blakeborough J. Cleveland bay horse, its history, evolution and importance today. 1950.

10. Reese HH. Breeds of light horses. US Department of Agriculture; 1918.

11. Russell D. The lives and legends of Buffalo Bill. University of Oklahoma Press; 1979.

12. Johnson D. Horse breeds. Voyageur Press; 2008.

13. Walling G. Cleveland Bay Horse Society Studbook Vol XXXIII. Clevel Bay Horse Soc. 1994.

14. Mills LS, Ballou JD, Gilpin M, Foose TJ. Population Management for Survival and Recovery: Analytical Methods and Strategies in Small Population Conservation. J Wildl Manage. 1997. doi:10.2307/3802439

15. Sargolzaei M, Iwaisaki H, Colleau JJ. Efficient computation of the inverse of gametic relationship matrix for a marked QTL. Genet Sel Evol. 2006. doi:10.1051/gse:2006002

16. Boichard D, Maignel L, Verrier É. The value of using probabilities of gene origin to measure genetic variability in a population. Genet Sel Evol. 1997. doi:10.1186/1297-9686-29-1-5

17. Groeneveld E, Westhuizen BD, Maiwashe A, Voordewind F, Ferraz JB. POPREP: a generic report for population management. Genet Mol Res. 2009. doi:10.4238/vol8-3gmr648

18. Chikhi L, Bruford M. Mammalian population genetics and genomics. Mammalian Genomics. 2004. doi:10.1079/9780851999104.0539

19. Cornuet JM, Luikart G. Description and power analysis of two tests for detecting recent population bottlenecks from allele frequency data. Genetics. 1996. doi:10.1093/oxfordjournals.jhered.a111627

20. Weir BS, Cockerham CC. Estimating F-Statistics for the Analysis of Population Structure. Evolution (N Y). 1984. doi:10.2307/2408641

21. Weir BS, Hill WG. Estimating F-Statistics. Annu Rev Genet. 2002. doi:10.1146/annurev.genet.36.050802.093940

22. Nei M. Molecular evolutionary genetics. Columbia university press; 1987.

23. Alix B, Boubacar DA, Vladimir M. T-REX: A web server for inferring, validating and visualizing phylogenetic trees and networks. Nucleic Acids Res. 2012. doi:10.1093/nar/gks485

24. Peakall R, Smouse PE. GenAlEx 6.5: genetic analysis in Excel. Population genetic software for teaching and research-an update. Bioinformatics. 2012;28: 2537–2539. doi:10.1093/bioinformatics/bts460

25. Huson DH, Bryant D. Application of phylogenetic networks in evolutionary studies. Molecular Biology and Evolution. 2006. doi:10.1093/molbev/msj030

26. Pritchard JK. Documentation for structure software : Version 2. 2. Statistics (Ber). 2007.

27. Corander J, Waldmann P, Marttinen P, Sillanpää MJ. BAPS 2: Enhanced possibilities for the analysis of genetic population structure. Bioinformatics. 2004. doi:10.1093/bioinformatics/bth250

28. Valera M, Molina A, Gutiérrez JP, Gómez J, Goyache F. Pedigree analysis in the Andalusian horse: Population structure, genetic variability and influence of the Carthusian strain. Livest Prod Sci. 2005. doi:10.1016/j.livprodsci.2004.12.004

29. Zechner P, Sölkner J, Bodo I, Druml T, Baumung R, Achmann R, et al. Analysis of diversity and population structure in the Lipizzan horse breed based on pedigree information. Livest Prod Sci. 2002. doi:10.1016/S0301-6226(02)00079-9

30. Gutiérrez JP, Marmi J, Goyache F, Jordana J. Pedigree information reveals moderate to high levels of inbreeding and a weak population structure in the endangered Catalonian donkey breed. J Anim Breed Genet. 2005. doi:10.1111/j.1439-0388.2005.00546.x

31. Aberle K, Wrede J, Distl O. Analyse der Populationsstruktur des Süddeutschen Kaltbluts in Bayern. Berl Munch Tierarztl Wochenschr. 2004.

32. Fox-Clipsham LY, Brown EE, Carter SD, Swinburne JE. Population screening of endangered horse breeds for the foal immunodeficiency syndrome mutation. Vet Rec. 2011. doi:10.1136/vr.100235

33. Prystupa JM, Hind P, Cothran EG, Plante Y. Maternal lineages in native canadian equine populations and their relationship to the nordic and mountain and moorland pony breeds. J Hered. 2012. doi:10.1093/jhered/ess003

34. McGahern AM, Edwards CJ, Bower MA, Heffernan A, Park SDE, Brophy PO, et al. Mitochondrial DNA sequence diversity in extant Irish horse populations and in ancient horses. Anim Genet. 2006. doi:10.1111/j.1365-2052.2006.01506.x

35. Brinkmann L, Gerken M, Riek A. Adaptation strategies to seasonal changes in environmental conditions of a domesticated horse breed, the Shetland pony (Equus ferus caballus). J Exp Biol. 2012. doi:10.1242/jeb.064832

36. Gu J, Orr N, Park SD, Katz LM, Sulimova G, MacHugh DE, et al. A genome scan for positive selection in thoroughbred horses. PLoS One. 2009. doi:10.1371/journal.pone.0005767

37. Khanshour A, Conant E, Juras R, Cothran EG. Microsatellite analysis of genetic diversity and population structure of Arabian horse populations. J Hered. 2013;104: 386–398.

38. Achmann R, Curik I, Dovc P, Kavar T, Bodo I, Habe F, et al. Microsatellite diversity, population subdivision and gene flow in the Lipizzan horse. Anim Genet. 2004. doi:10.1111/j.1365-2052.2004.01157.x

39. Luís C, Juras R, Oom MM, Cothran EG. Genetic diversity and relationships of Portuguese and other horse breeds based on protein and microsatellite loci variation. Anim Genet. 2007. doi:10.1111/j.1365-2052.2006.01545.x

40. Cañon J, Checa ML, Carleos C, Vega-Pla JL, Vallejo M, Dunner S. The genetic structure of Spanish Celtic horse breeds inferred from microsatellite data. Anim Genet. 2000. doi:10.1046/j.1365-2052.2000.00591.x

41. Juras R, Cothran EG, Klimas R. Genetic Analysis of Three Lithuanian Native Horse Breeds. Acta Agric Scand - Sect A Anim Sci. 2003. doi:10.1080/09064700310012971

42. Luís C, Cothran EG, Oom MDM. Inbreeding and genetic structure in the endangered Sorraia horse breed: Implications for its conservation and management. J Hered. 2007. doi:10.1093/jhered/esm009

43. Luikart G, Cornuet JM. Empirical evaluation of a test for identifying recently bottlenecked populations from allele frequency data. Conserv Biol. 1998. doi:10.1046/j.1523-1739.1998.96388.x

44. Lacy RC. Should we select genetic alleles in our conservation breeding programs? Zoo Biology. 2000. doi:10.1002/1098-2361(2000)19:4<279::AID-ZOO5>3.0.CO;2-V

45. Pearl MC. Research Techniques in Animal Ecology Methods and Cases in Conservation Science. J Wildl Manage. 2000. doi:10.2307/3803113

46. Fraser DJ, Bernatchez L. Adaptive evolutionary conservation: Towards a unified concept for defining conservation units. Molecular Ecology. 2001. doi:10.1046/j.1365-294X.2001.t01-1-01411.x

47. Lacy RC. Management of limited animal populations. Bottlenose dolphin reproduction workshop. 2000.

48. Ansari-Mahyari S, Berg P. Power of QTL mapping using both phenotype and genotype information in selective genotyping.

49. Maccluer JW, Boyce AJ, Dyke B, Weitkamp LR, Pfenning DW, Parsons CJ. Inbreeding and pedigree structure in standardbred horses. J Hered. 1983. doi:10.1093/oxfordjournals.jhered.a109824

50. G. J-M, Falconer DS. Introduction to Quantitative Genetics. Popul (French Ed. 1962. doi:10.2307/1525780

51. Gutiérrez JP, Goyache F. A note on ENDOG: A computer program for analysing pedigree information. J Anim Breed Genet. 2005. doi:10.1111/j.1439-0388.2005.00512.x

52. Lacy RC. Analysis of founder representation in pedigrees: Founder equivalents and founder genome equivalents. Zoo Biol. 1989. doi:10.1002/zoo.1430080203

53. Bijma P, Woolliams JA. Prediction of genetic contributions and generation intervals in populations with overlapping generations under selection. Genetics. 1999.

54. Sørensen AC, Sørensen MK, Berg P. Inbreeding in danish dairy cattle breeds. J Dairy Sci. 2005. doi:10.3168/jds.S0022-0302(05)72861-7

55. Lacy RC. Clarification of genetic terms and their use in the management of captive populations. Zoo Biol. 1995. doi:10.1002/zoo.1430140609

56. Belkhir K, Borsa P, Chikhi L, Raufaste N, Bonhomme F. GENETIX 4.05, Windows TM software for population genetics. Lab génome, Popul Interact CNRS Umr. 2004;5000.

57. Hill WG. Molecular Evolutionary Genetics. By Masatoshi Nei. New York: Columbia University Press. 1987. 512 pages. U.S. $50.00. ISBN 0 231 06320 2. Genet Res. 1988. doi:10.1017/s001667230002735x

58. Wright S. Variability within and among natural populations. Evolution and the genetics of populations. 1978.

59. Caballero A, Toro MA. Analysis of genetic diversity for the management of conserved subdivided populations. Conserv Genet. 2002. doi:10.1023/A:1019956205473

60. Makarenkov V. T-REX: Reconstructing and visualizing phylogenetic trees and reticulation networks. Bioinformatics. 2001. doi:10.1093/bioinformatics/17.7.664

